# Divergent evolutionary strategies preempt tissue collision in fly gastrulation

**DOI:** 10.1101/2023.10.09.561568

**Authors:** Bipasha Dey, Verena Kaul, Girish Kale, Maily Scorcelletti, Michiko Takeda, Yu-Chiun Wang, Steffen Lemke

## Abstract

Metazoan development proceeds through a series of morphogenetic events that sculpt body plans and organ structures. In the early embryonic stages, morphogenetic processes involving growth and deformation occur concurrently. Forces generated in one tissue can thus increase mechanical stress in the neighboring tissue, potentially disrupting spatial patterning, morphological robustness, and consequently decreasing organismal fitness. How organisms evolved mechanisms to reduce or release inter-tissue stresses remains poorly understood. Here we combined phylogenetic survey across a whole insect order (Diptera), quantitative live imaging, and functional mechanical perturbation to investigate the evolution of mechanical stress management during epithelial expansions in the gastrulating fly embryos. We find that two distinct cellular mechanisms exist in Diptera to prevent the accumulation of compressive stress that can arise when the expanding head and trunk tissues collide. In Cyclorrhapha, a monophyletic dipteran subgroup including the fruit fly *Drosophila melanogaster*, the head-trunk boundary undergoes active out-of-plane deformation to form a transient epithelial fold, called the cephalic furrow (CF), which acts as a mechanical sink to preempt head-trunk collision. Genetic or optogenetic elimination of the CF leads to tissue buckling, yielding deleterious effects of axial distortion that likely results from unmitigated release of compressive stress. Non-cyclorrhaphan flies, by contrast, lack CF formation and instead display widespread out-of-plane division in the head, which shortens the duration of its expansion and reduces surface area increase. Reorienting head mitosis in *Drosophila* from in-plane to out-of-plane partially suppresses the need for epithelial out-of-plane deformation, suggesting that out-of-plane division can act as an alternative mechanical sink to prevent tissue collision. Our data suggest that programs of mechanical stress management can emerge abruptly under selective pressure of inter-tissue mechanical conflict in early embryonic development.

## Main text

The development of multicellular organisms proceeds through a series of morphogenetic events that sculpt tissue morphology. Morphogenesis is fundamentally a mechanical process that involves both tissue-intrinsic active stress as well as passive deformation caused by external forces. In animals, morphogenesis starts during gastrulation, when simple cell clusters or sheets are transformed into more complex embryonic tissues with inner layers and curved shapes. During gastrulation, morphogenetic processes occur simultaneously within a mechanical continuum, characterized by the absence of clear spatial segregation and distinct material compartmentalization. This raises the question of how forces emanating from one morphogenetic process may influence another. One possibility is that extrinsic forces may be co-opted to facilitate local deformation^1–6^. In cases where conflict between adjacent tissues leads to the accumulation of mechanical stress, however, it is unclear whether the orderedness and robustness of development is contingent upon the evolution of specific mechanisms capable of managing the stress. Very few studies have examined how organisms deal with inter-tissue morphogenetic stress. Revealing the evolutionary trajectory of mechanical stress management would require: one, a combined effort of well-resolved phylogeny; two, broad sampling and documentation of phenotypic states from representative clades; and three, functional assessment through genetic and mechanical perturbation.

Gastrulation in the fruit flies *Drosophila melanogaster* (henceforth *Drosophila*) begins with concurrent morphogenetic movements of all three germ layers^7,8^. At gastrulation onset, mesoderm internalizes through ventral furrow (VF) formation, endoderm internalization begins with posterior midgut (PMG) invagination, and in the region where the head ectoderm abuts the trunk ectoderm forms a deep epithelial fold, called the cephalic furrow (CF)^9–11^. The onset of these morphogenetic movements are immediately followed by a convergent extension event called germband extension (GBE) that elongates the anterior-posterior (A-P) axis of the trunk ectoderm^12^. Coincident with GBE, a series of locally synchronous events of mitosis, localized to the so-called mitotic domains (MDs), occur in a spatially stereotypical and temporally ordered manner, with 4 out of 5 of them located in the head ectoderm^13^. Thus, the first 30 min of *Drosophila* gastrulation is characterized by the simultaneous occurrence of multiple morphogenetic events. Among these, the CF is unique. It is a transient epithelial fold that forms and retracts back to the embryonic surface, but does not give rise to any internal cell type or tissue structure^13^. Because CF positioning is precise^9,14^, it is likely that its formation is critical for embryonic development, although its function remains unknown. One hypothesis, based on the fact that the CF is formed specifically at the head-trunk boundary, proposes that the CF functions as an anterior barrier to guide the long-range, posterior-directed flow of GBE^15^. This hypothesis would predict that the CF is unique to insects that undergo GBE. A contrasting hypothesis posits deep conservation between the CF and the vertebrate head-trunk boundary on the basis of homologous gene expression^16^. Neither hypothesis has as of yet been put to rigorous tests of phylogenetic and functional analysis.

## The cephalic furrow: a morphogenetic innovation at the head-trunk interface of Cyclorrhaphan flies

To gain insights into the evolutionary history and the developmental function of the CF, we conducted a phylogenetic survey and asked whether the CF is universal throughout the insect order of Diptera. We comprehensively surveyed classic and modern literature (Fig. 1 Supplementary table 1), and complemented that with our own imaging data and analyses in species that are phylogenetically informative. We defined the CF as a deep epithelial fold formed at the head-trunk boundary in the early gastrula and found that it is present only in embryos of cyclorrhaphan flies, but not in non-cyclorrhaphan flies (Fig. 1a-f; Fig. 1 Supplement 1). In contrast, there is clear evidence for GBE in all dipteran species that we have examined or for which we could find reference images (Fig. 1 Supplement 1; Fig. 1 Supplementary table 1). These data suggest that the CF is an evolutionary novelty and a synapomorphic character for the monophyletic group of Cyclorrhapha.

**Fig. 1.**
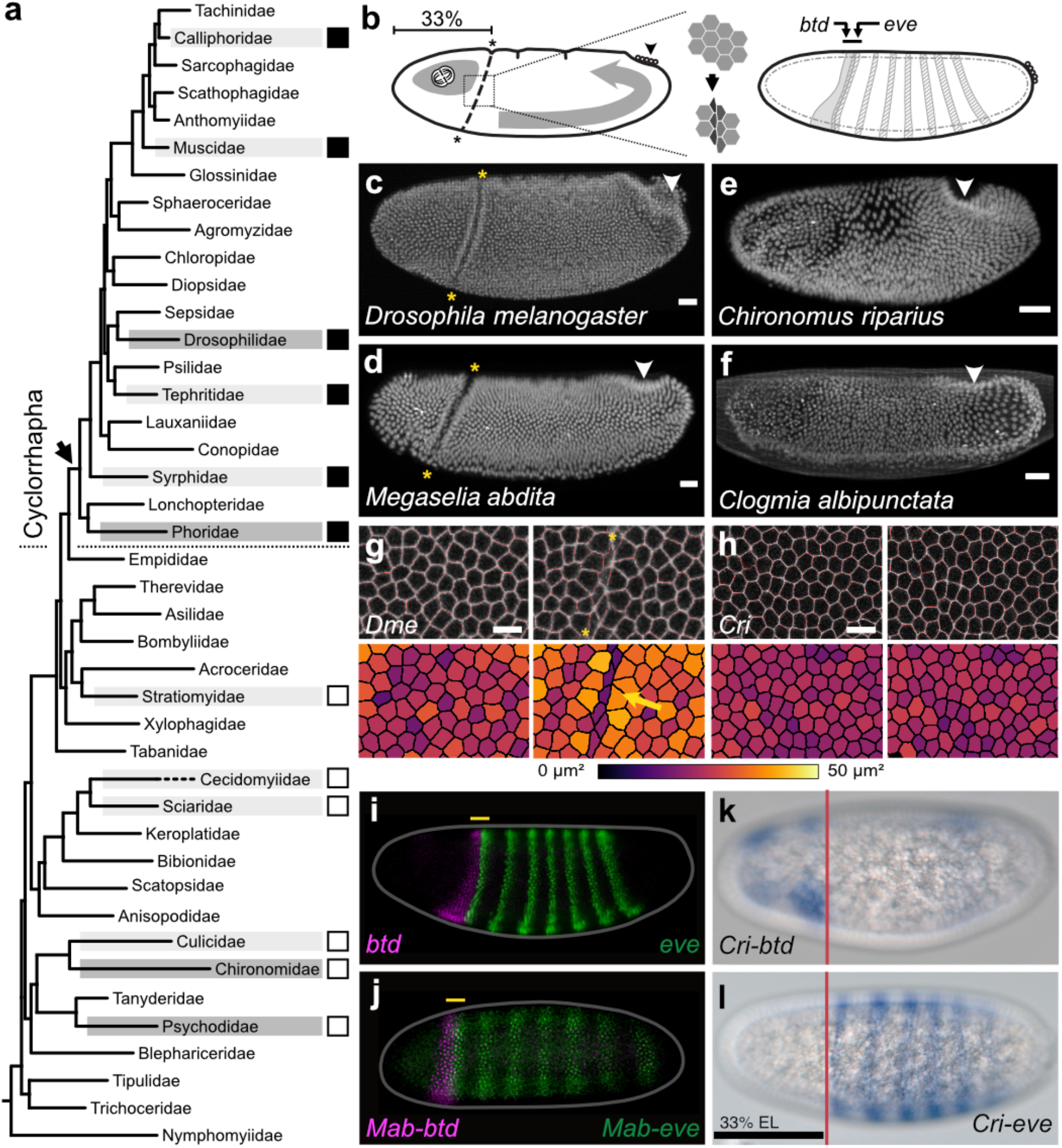
| The CF is an evolutionary innovation at the head-trunk interface of flies. **a**, Phylogenetic tree showing major fly families (adapted from Wiegmann et al., 2011^17^) marking the presence (filled boxes) or absence (empty boxes) of a CF in embryos belonging to various fly species, indicating that the CF appeared once (arrow) in the phylogeny of Diptera, and then did not disappear. Highlights indicate families for which species were evaluated (light) and studied (dark) in this work. See also, Fig. 1 Supplementary table 1. **b**, The schematics summarize our understanding of the spatio-temporal occurrence of the CF (its location in embryo coordinates and its chronology relative to other morphogenetic events), the causal tissue level changes, and the genetic program that dictates its formation. **c-f**, Representative images of fixed embryos at comparable stages from the four selected species, *Drosophila melanogaster* (Drosophilidae) (**c**, n=13 embryos), *Megaselia abdita* (Phoridae) (**d**, n=13 embryos), *Chironomus riparius* (Chironomidae) (**e**, n=24 embryos), and *Clogmia albipunctata* (Psychodidae) (**f**, n=13 embryos). White arrowheads, the extent of ectoderm elongation; yellow asterisks, the CF. Scale bars, 25 µm. **g**, **h**, Mesoscopic view of the head-trunk boundary region in *Drosophila* (**g**, n=3) and *Chironomus* (**h**, n=3) embryos at t=-3 (left) or 0 (right) min relative to the onset of gastrulation. Asterisks, the CF region; arrow, the reduction in apical cell areas in *Drosophila*. Such reduction in apical area is not observed in *Chironomus* embryos. LUT bar, the color-code used for cell area. Scale bars, 10 µm. **i**, **j**, Maximum projections showing the partial overlap (yellow line) of Btd (red) and Eve (green) expression patterns via immunofluorescence in *Drosophila* (**i**, n=3 embryos) or fluorescent *in situ* hybridization for *Mab-btd* and *Mab-eve* in *Megaselia* (**j**, n=3 embryos). **k, l**, *in-situ* hybridization reveals transcript distributions of *Cri-btd* (**k**, n=3 embryos) and *Cri-eve* (**l**, n=3 embryos) in *Chironomus*; stainings are aligned along the A-P axis and indicate a lack of overlap in their expression patterns. Vertical lines, the anterior boundary of *Cri-eve1* at 33%EL (egg length).

Direct comparison between *Drosophila* and the midge *Chironomus riparius* (henceforth *Chironomus*), a representative non-cyclorrhaphan species (Fig. 1a; Chironomidae), provides the most unequivocal evidence yet that non-cyclorrhaphan flies do not form the CF. In *Drosophila*, CF initiation is concurrent with the onset of PMG invagination^9^, and the furrow structure persists for about 90 min before it fully retracts back to the surface. Live imaging of *Chironomus* embryos reveals a complete lack of infoldings at the head-trunk boundary, from the blastoderm stage to the end of GBE (Fig. 1 Supplement 1). In *Drosophila*, CF initiation occurs at ∼33% embryo length (EL) where the apical cell surface area can be seen to decrease in one to two columns of linearly aligned initiating cells^9,14^. To visualize this, we color-coded cell apical surface areas in the region between ∼28 to 39% EL (Fig. 1g). In *Chironomus*, no such apical surface decrease could be observed in the area between ∼28 to 52% EL (Fig. 1h). Furthermore, while we could observe progressive internalization of cells into the CF in *Drosophila*, incorporating cells from both the head and the trunk ectoderm (Fig. 1 Supplement 2a, and Fig. 1 Movie S1 top panel, both show ∼10 to 75% EL), no such cell internalization could be seen at the head-trunk boundary in *Chironomus* (Fig. 1 Supplement 2b, and Fig. 1 Movie S1 bottom panel, both show ∼15 to 85% EL). We conclude that *Chironomus* embryos do not form CF.

Finally, CF formation in *Drosophila* requires overlapping expression of the transcription factors Buttonhead (Btd) and Even-skipped (Eve), which combinatorially define a single or a double column of cells where the lateral membrane contracts to initiate the CF^9,10^ (Fig. 1i). *Megaselia abdita* (henceforth, *Megaselia*), a representative species that branched off near the cyclorrhaphan stem group (Fig. 1a, Phoridae)^17^, forms the CF (Fig. 1d) and shows such an overlap (Fig. 1j). This suggests the Btd/Eve overlap is a conserved feature associated with CF formation in Cyclorrhapha, and predicts its absence in non-cyclorrhaphan flies. To test this, we cloned the *Chironomus* orthologs of *btd* and *eve*, and analyzed their expression pattern (*Cri-btd* and *Cri-eve*, see Fig. 1 Supplement 3 for the protein tree). In blastoderm stage embryos, *Cri-btd* is expressed in two separate domains in the head region (Fig. 1k); *Cri-eve* is expressed in six stripes, while the seventh pair-rule stripe comes up at the posterior pole after the onset of GBE (Fig. 1l). Importantly, the patterns do not overlap (Fig. 1k, l). These data suggest non-cyclorrhaphan flies lack the positional code necessary for CF initiation. Indeed, another non-cyclorrhaphan fly *Clogmia albipunctata* (henceforth *Clogmia*; Fig. 1a, Psychodidae) lacks the CF (Fig. 1f; Fig. 1 Supplement 1), and, as demonstrated in an independent study by Vellutini *et al.* ^(footnote 1)^, shows no overlapping expression between *btd* and *eve*^18^. These results suggest that the evolutionary origin of the CF is associated with a change of the expression domain of *btd* and the gain of its overlap with the first stripe of *eve* (called *eve1* hereafter) in the stem group of Cyclorrhapha. We conclude that the CF is a morphogenetic innovation with a complex underlying program of genetic patterning, cellular mechanics and tissue behaviors, none of which could be detected in *Chironomus*, and are expected to be absent in all other non-cyclorrhaphan flies.

## Genetic and optogenetic ablation of CF causes ectodermal buckling at the head-trunk interface

To elucidate how the CF arose as an evolutionary novelty, we sought to characterize the functional role of the CF during development. This has been challenging because both *btd* and *eve* mutants lead to significant patterning defects outside the head-trunk boundary. To overcome this, we blocked CF formation by specifically eliminating the expression of *eve1*. We engineered a full-length *eve* genomic construct, removing enhancer elements that confer the expression of *eve1*, while leaving all other cis-regulatory elements intact^19,20^ (Fig. 2 Supplement 1a; see Methods). We introduced this into an *eve* null genetic background, yielding the *eve1^KO^* line, and confirmed that the *eve1^KO^* embryos lack *eve1* expression (Fig. 2 Supplement 1b). We then performed live imaging and found that at gastrulation onset they lack planar polarized Myosin accumulation in the cells predicted to initiate the CF^9^ and consequently do not form the CF (Fig. 2a, b; Fig. 2 Movie S1). Instead, the head-trunk interface undergoes a late stage out-of-plane deformation (Fig. 2c, d; Fig. 2 Movie S2) referred to as the ‘head-trunk buckles’ hereafter for the following reasons. First, they occur without localized Myosin enrichment (Fig. 2a, b; Fig. 2 Movie S1). Second, cells that initiate the head-trunk buckle move inward abruptly, rather than undergoing gradual shortening as seen in a normal CF, leading to a large surface indentation that differs from the narrow surface cleft of CF (Fig. 2a, b; Fig. 2 Movie S1). Third, the head-trunk buckle occurs ∼12 min after the onset of PMG invagination, whereas CF initiation and PMG onset are concurrent (Fig. 2i; Fig. 2 Movie S2). Lastly, the D-V position of the head-trunk buckles vary among embryos (Fig. 2 Supplement 1c), contrasting with the CF, which typically starts laterally and spreads dorsally and ventrally^9^. The lack of localized Myosin, the lack of a stereotypical spatial pattern, the delayed timing, and the abruptness with which the buckles are formed without localized cell shape change, all suggest that these are not genetically patterned active deformations, but are consistent with passive buckling. Similar head-trunk buckles could also be observed in the classic *eve* and *btd* mutants (Fig. 2e, f, i; Fig. 2 Movie S2), confirming previous reports^9,10^ and the independent study by Vellutini *et al.*^18^, thus validating the use of *eve* or *btd* mutants, or global RNAi knockdown (see below), in generating the head-trunk buckles.

**Fig. 2.**
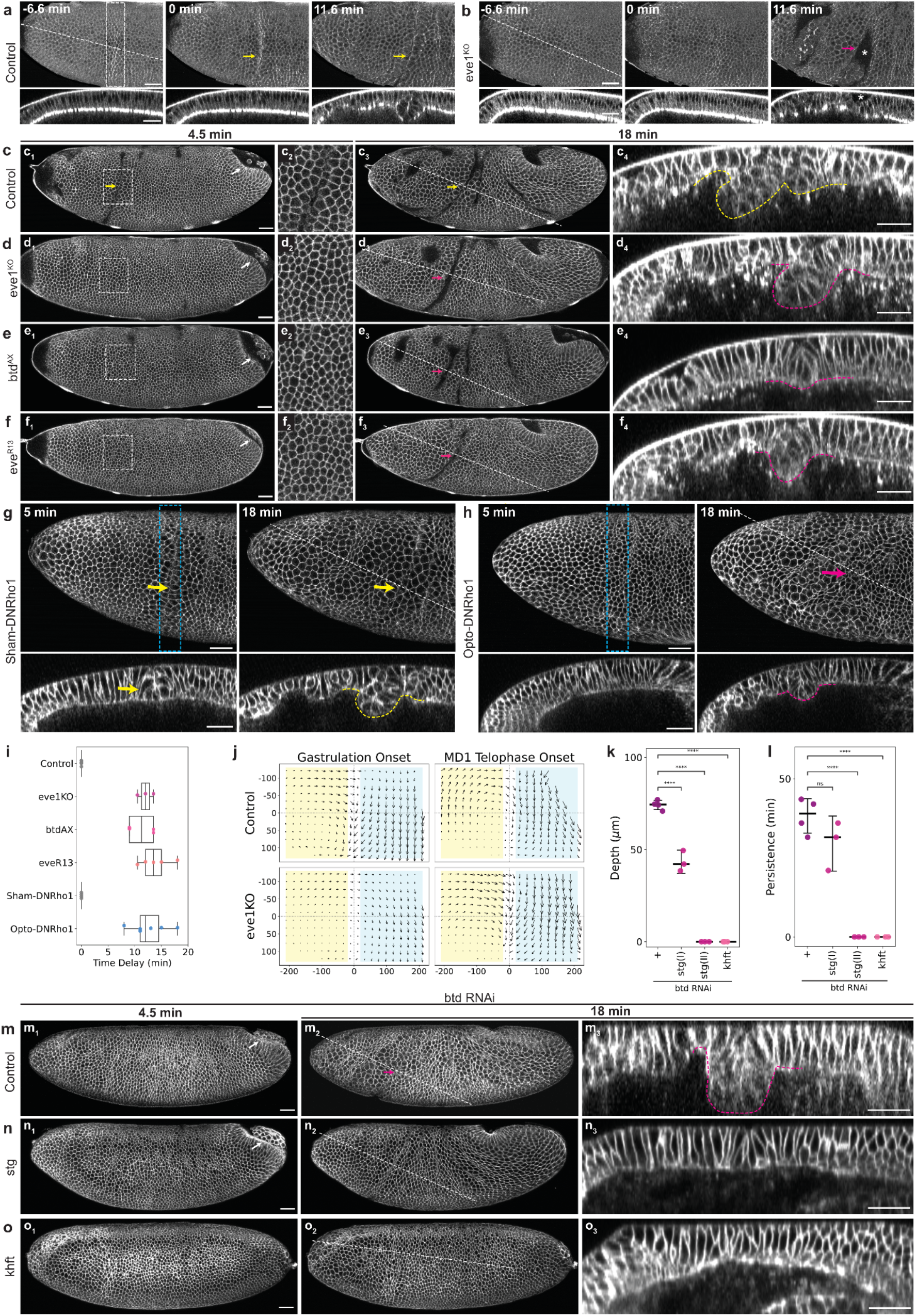
| The CF functions as a mechanical sink to prevent tissue collision. **a**, **b**, Time-lapse series of a representative control (**a**, *eve1^KO^* /+, n=3) or *eve1^KO^* (**b**, n=4) embryo expressing MyoII-mKate2 showing a lateral surface projection (top rows) of the anterior half of the embryo that includes the head-trunk boundary and a z-axis reslice (bottom rows). The z-axis reslices were generated along the white dashed lines drawn on the lateral surface projection view. Note the reslice views were cropped such that the lengths of the dashed line do not correspond to the widths of the reslice view. This style of annotation and reslice representation applies to the reslices in all figures. Dashed rectangle denotes planar polarized MyoII accumulation prior to CF initiation; asterisks, wide surface indentation resulting from head-trunk buckling. **c**–**f**, Time-lapse series of a representative control (**c**, *Gap43-mCherry/+*, n=6), *eve1^KO^*(**d**, n=3), *btd^AX^* (**e**, n=4), or *eve^R13^* (**f**, n=6) embryo, visualized with a membrane marker (Gap43-mCherry) showing a lateral surface projection (c1-f1; c3-f3), an enlarged view of the CF region (c2-f2, of the area highlighted with a dashed rectangle in c1-f1) and a z-axis reslice (c4-f4). **g**, **h**, Time-lapse series of a representative sham control (**g**, Sham-DNRho1, n=4) or a photoactivated (**h**, Opto-DNRho1, n=7) embryo expressing the Opto-DNRho1 system, visualized with a membrane marker (3xmScarlet-CaaX) showing a lateral surface projection of the anterior half of the embryo that includes the head-trunk boundary (top rows) and a z-axis reslice (bottom rows). Blue dashed rectangles, illumination ROIs for the sham activation or photoactivation. **i**, Timing of the onset of CF or head-trunk buckling relative to gastrulation onset. Sample size, Control=6, *eve1^KO^*=3, *btd^AX^* =4, *eve^R13^=5*, Sham-DNRho1=3, Opto-DNRho1=6. **j**, Tissue flow fields visualized with PIV in control (n=3) and *eve1^KO^* (n=2) at the onset of gastrulation and MD1 telophase. Yellow shaded rectangles, head region; blue shaded rectangles, trunk regions; x-origin, head-trunk boundary; y-origin, lateral midline. Unit, pixel. **k**, **l**, Maximum depths (**k**) or durations (**l**) of head-trunk buckles in *btd* RNAi embryos with additional genetic manipulations plotted with median and error bars indicating 95% confidence interval. The *stg* mutants show two classes (I and II) of phenotypes. One-way ANOVA Tukey post-hoc test; ****, p < 0.0001; ns, not significant (p > 0.05). Sample size: Control=4, *stg*(I)=3, *stg*(II)=3, *khft*=3. **m**, **n**, **o**, Time-lapse series of *btd* RNAi embryos in control (**m**, n=5), *stg* (**n**, n=6), or *khft* (**o**, n=3) genetic background, visualized with a membrane marker (3xmScarlet-CaaX) showing a lateral surface projection (m1-o1; m2-o2) and a z-axis reslice (m3-o3). Sample size, same as in **k**, **l**, Common annotations: white arrows, PMG; yellow arrows and yellow dashed outlines, CFs; magenta arrows and magenta dashed outlines, head-trunk buckles. Time is relative to the onset of gastrulation. Scale bars, 30 µm.

To rule out the possibility that the head-trunk buckles arise from local alteration of genetic patterning, we mechanically blocked CF formation using an optogenetic system, the opto-DNRho1 system^9,21^, that inhibits actomyosin contractility. To block CF formation, we illuminated the entire CF region on one side of the embryo with a narrow ROI (See Methods) so as not to perturb contractility elsewhere. This treatment completely eliminates CF formation, and buckling was observed at the head-trunk boundary similar to the *eve1^KO^* embryo (Fig. 2g-i; Fig. 2 Movie S3). This reaffirms our assertion that the head-trunk interface buckles in the absence of CF and that buckling occurs in the absence of local increase of actomyosin contractility given the presence of photoactivated opto-DNRho1. It also rules out local genetic perturbation as a probable cause of buckling as the opto-DNRho1 system does not incur patterning change. Thus, both genetic and mechanical blockage of CF initiation results in passive buckling, raising the possibility that when the CF does not form, compressive stress accumulates in the ectodermal regions flanking the head-trunk interface.

In search of the potential source of compressive stress, we made two observations. First, we observed a correlation between mitosis and buckling in embryos in which the CF is optogenetically inhibited, as the head-trunk buckle typically starts when the cells of the MD2, the second mitotic domain, undergo mitotic rounding (Fig. 2 Supplement 1d; Fig. 2 Movie S3). This also confirms that buckling is delayed relative to the normal timing of CF initiation, as in the wild-type the MD2 cells round up well after CF initiation, when a substantial furrow structure is already present (Fig. 2 Supplement 1d; Fig. 2 Movie S3). Second, the anterior edge of the trunk ectoderm can be seen as moving inward as the buckle deepens out-of-plane (Fig. 2 Movie S3), suggesting that GBE may be involved in the head-trunk buckling. In sum, the head-trunk buckling could be related to compressive stress generated during head expansion via mitosis and trunk expansion via convergent-extension.

## Collision of two persistent, coherent tissue flows at the head-trunk interface likely leads to accumulation of compressive stress

To gain insights into how compressive stress arises at the head-trunk boundary, we characterized the tissue flow field using particle image velocimetry (PIV) and analyzed it relative to the head-trunk boundary and the lateral midline (See Methods and Fig. 2 Supplement 1e for a representative image of PIV superimposed on the embryo image.). In the wild-type embryos, we observed a local, convergent flow at the head-trunk boundary where the CF is formed, suggesting that the CF behaves as a tissue sink and that the head and trunk tissues flow into this sink (Fig. 2j, control at gastrulation onset). Indeed, ∼5 columns of cells on both the head and trunk sides adjacent to the CF initiating cells become incorporated into the CF (Fig. 1 Supplement 2a; Fig. 1 Movie S1 top panel). PIV analysis also reveals that the CF breaks the continuity of the flow field by separating it into two persistent, regionally coherent flows in the flanking tissues – a posterior-ward flow in the head and a ventral-ward flow in the trunk that diverges along the A-P axis (Fig. 2j, control at MD1 telophase onset). This suggests that without a sink, the head and the trunk tissue flows would have collided at the head-trunk interface. In contrast, the *eve1^KO^* embryo displays a single, uninterrupted flow field at the onset of gastrulation, as predicted in a previous computational model when the CF is absent^15^ (Fig. 2j, *eve1^KO^*at gastrulation onset). This supports the premise that the CF functions as a sink. As the embryo begins to buckle, a local convergent flow emerges at the head-trunk boundary, similar to the one we observed at the CF in the wild-type, suggesting that the buckle also behaves as a sink (Fig. 2j, *eve1^KO^* at MD1 telophase onset). The head-trunk buckle breaks the continuity of the flow field, similar to the CF, further supporting the idea that the head-trunk boundary interfaces two colliding tissue flows. We observed similar flow fields in embryos that lack Btd expression, confirming these analyses (Fig. 2 Supplement 1f). Analogous to the collision of tectonic plates, which can lead to out-of-plane deformation of Earth’s crust, we propose that tissue flows driven by the expanding head and trunk regions undergo ‘tissue tectonic collision’ at the head-trunk boundary, causing local accumulation of compressive stress and leading to tissue buckling when the genetically programmed sink, the CF, is absent.

## The depth and persistence of head-trunk buckle is reduced when mitosis or trunk convergent extension is blocked

To test this hypothesis, we asked whether a reduction of tissue flow in either the head or trunk dampens buckling. In case of head expansion, we consider the possibility that cell division in the head MDs drives the posterior-ward tissue flow. As the columnar epithelial cells divide, they round up during metaphase and transiently expand their apical surface (see below). Such a surface expansion can effectively enlarge the entire head region, leading to the posterior-ward tissue flow, because the embryo is bilaterally symmetric and spatially confined within the stiff vitelline membrane. To block cell division, we removed the zygotic activity of String (Stg)/Cdc25, which activates Cdk1 to drive mitosis in each MD^22,23^. In *stg/cdc25* mutants, CF initiation occurs normally, indicating that mitosis in the head MDs is not required for CF initiation (Fig. 2 Supplement 2a, c, d; Fig. 2 Movie S4). To examine the effect of division-driven head expansion on tissue collision at the head-trunk boundary, we induced buckling using RNAi knockdown of *btd* in *stg/cdc25* mutants. We observed two classes of phenotype: class I exhibits late onset of buckling with reduced depth and persistence (*stg(I)*; Fig. 2k, l; Fig. 2 Supplement 2b, e; Fig. 2 Movies S4, S5), while class II shows complete lack of buckling (*stg(II)*; Fig. 2k-n), similar to the phenotype reported in the independent study by Vellutini *et al.*^18^. These data thus support the hypothesis that head expansion contributes to buckling, presumably by driving the posterior-ward flow of the head that collides with the trunk.

In the trunk, tissue flow stems from the combined effect of VF formation and GBE^15^the former of which is known to drive the ventral-ward flow^15^, while the latter likely accounts for the A-P divergent flow, in light of our PIV analysis that shows the anterior-ward flow of the trunk ectoderm (Fig. 2j), in addition to the well-characterized posterior-ward flow^2^. To examine whether reduction of trunk extension can dampen buckling, we used a quadruple mutant line of *knirps hunchback forkhead* and *tailless* (*khft*) to abrogate GBE^24,25^. The *khft* quadruple mutant eliminates both local junctional transition and external drag force exerted by PMG invagination, the two active processes involved in GBE^2^, and thus its trunk ectoderm exhibits no axis elongation whatsoever (Fig. 2 Supplement 2a; Fig. 2 Movie S4). In contrast, CF initiation is normal, indicating that CF initiation does not require GBE. We then inhibited CF initiation using *btd* RNAi in the *khft* mutant and saw no head-trunk buckling at all (Fig. 2k-m, o; but note the existence of late buckling linked to mitoses, Fig. 2 Supplement 2b, e; Fig. 2 Movies S4, S5). These data suggest that trunk extension along the A-P axis contributes to buckling, likely due to a GBE-driven, A-P divergent flow that collides with the head ectoderm.

Together, data presented above support our hypothesis that genetically programmed tissue expansion in the head and trunk results in tissue collision, producing head-trunk buckles when the CF does not form. The expanding head and trunk indeed ‘fuel’ the CF, as both *stg* and *khft* mutants form shallower and less persistent CF than the wild-type (Fig. 2 Supplement 2c, d). Thus, although the CF is initiated by the local increase of actomyosin contractility^9^, its subsequent, extensive invagination requires the expansion of the neighboring tissues. Given the pliability of invagination depth and persistence, we conclude that the CF has the capacity to ‘absorb’ the tissue surfaces of the expanding neighbors, acting as a *bona fide*, genetically patterned mechanical sink that guides the tissue flow, thereby preemptively preventing tissue collision and buckling at the head-trunk interface. This interpretation is further corroborated by the *in silico* simulation performed in the independent study by Vellutini *et al.*^18^.

## Abrogation of CF formation increases frequency of midline distortion

The precision and robustness with which the CF is formed suggest that its spatial patterning and temporal dynamics are under selective pressure. Thus, CF formation must confer fitness^9,14^. To test this, we asked whether head-trunk buckling has a deleterious effect on embryonic development. We eliminated the CF bilaterally using the opto-DNRho1 system such that the phenotypic effects can only be attributed to loss of the CF, but not altered genetic patterning. We then imaged the ventral half of the embryo to monitor embryonic development for ∼1.5 hours after the onset of gastrulation. We first confirmed that optogenetic perturbation indeed eliminates CF initiation as gastrulation commences (Fig. 3a, b; Fig. 3 Movie S1). The VF initiates normally, indicating the effect of optogenetic perturbation is restricted to the CF. VF formation continues with closure as the mesectodermal cells meet, resulting in a straight ventral midline. After the ventral midline forms, strikingly, we observed an increased frequency of midline distortion or rotation in embryos in which the CF is blocked, as compared to the sham control that maintains a bilaterally symmetric body plan (Fig. 3a, b, e; Fig. 3 Movie S1). In embryos in which the CF is optogenetically eliminated, the extent to which the ventral midline becomes distorted is variable, with strong midline distortion often associated with bilateral asymmetry of head-trunk buckling. These results suggest that releasing compressive stress via buckling is intrinsically stochastic, whereas programmed, active deformation, such the CF, reduces such stochasticity. We further confirmed that *btd* mutants (Fig. 3c, d, f; Fig. 3 Movie S2) and *btd* RNAi embryos (Fig. 3 Supplement 1) also show midline distortions. These data provide direct evidence that loss of the CF has a deleterious effect on embryonic development that can be observed soon after gastrulation onset.

**Fig. 3.**
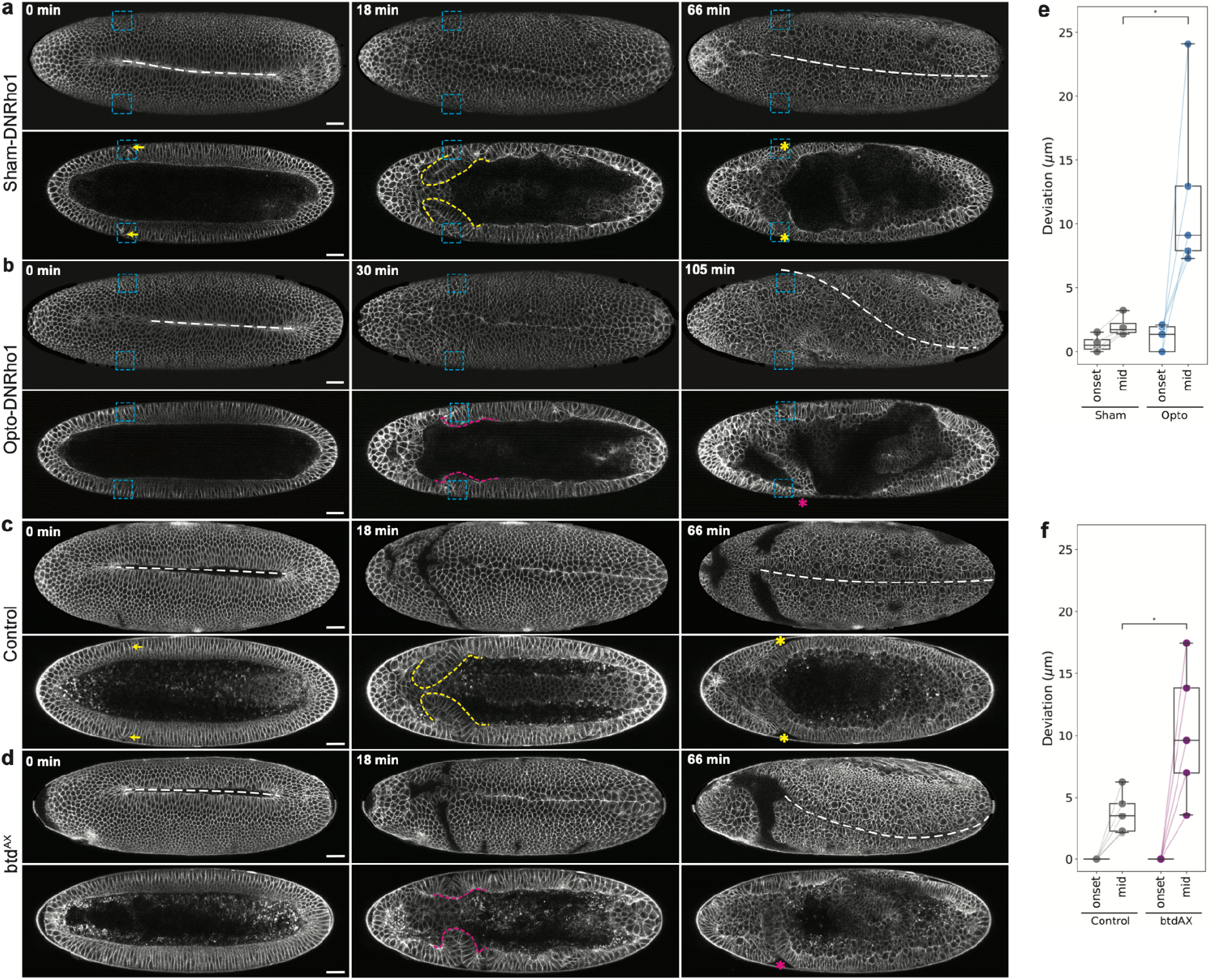
| Loss of the CF causes midline distortion. **a**–**d**, Time-lapse series of a representative sham control (**a**, Sham-DNRho1, n=4), or photoactivated (b, Opto-DNRho1, n=5) embryo expressing the Opto-DNRho1 system, visualized with 3xmScarlet-CaaX, or a control (**c**, *Gap43-mCherry/+*, n=6) or *btd^AX^*(**d**, n=6) embryo, visualized with Gap43-mCherry, showing a ventral surface projection (top rows) and a single coronal section (bottom rows). Blue dashed rectangles, illumination ROIs for the sham activation or photoactivation; white dashed lines, ventral midlines; yellow arrows and yellow dashed outlines, CF; yellow asterisks, bilaterally symmetric CFs; magenta arrows and magenta dashed outlines, head-trunk buckles; magenta asterisks, laterally asymmetric buckles. Time is relative to the onset of gastrulation. Scale bars, 30 µm. **e**, **f**, Box plots showing mean deviation of the manually-marked ventral midline from the expected linear ventral midline position measured at the onset of gastrulation (onset) and a mid-gastrulation (mid) stage when deviation reaches a maximum. Whiskers, min and max. Mann–Whitney U test; *, p < 0.05. Sample size: Sham=4, Opto=5, Control=4, *btd^AX^* =5.

## Mitosis in the head ectoderm is primarily oriented out-of-plane in non-Cyclorrhaphan flies

Our data thus far suggest that cyclorrhaphan flies, exemplified by *Drosophila*, avoid tissue collision at the head-trunk boundary and the accumulation of compressive stress via genetically programmed out-of-plane deformation in the form of the CF. Lacking the CF, the non-cyclorrhaphan embryos likely would need an alternative mechanism to dissipate compressive stress, since we find that cell density is not lower in the non-cyclorrhaphan embryos than in *Drosophila* (Fig. 4 Supplement 1), and that the trunk ectoderm undergoes axial elongation via GBE (Fig. 1 Supplement 1). We considered the possibility that early head morphogenesis differs between cyclorrhaphan and non-cyclorrhaphan flies. Using nuclei as a proxy for cells, we examined the epithelial morphology in the head region, defined as the anterior third of the embryo. We observed a monolayer epithelial architecture in the cyclorrhaphan flies *Drosophila* and *Megaselia* during early gastrulation (Fig. 4a). In contrast, in the non-cyclorrhaphan *Clogmia* and *Chironomus* a large area of the head epithelium displays a double layer of nuclei (Fig. 4a). Such double layering could result from pseudo-stratification or cell extrusion. We ruled these out through live imaging of the *Chironomus* embryos with a membrane marker (Fig. 4 Supplement 2). Instead, live imaging revealed that many mitotic cells form their cytokinetic ring parallel to the embryo surface, indicative of out-of-plane division (Fig. 4d). These data reveal that non-cyclorrhaphan flies differ from cyclorrhaphan flies in how cell divisions are oriented in the head region, which might affect head expansion.

**Fig. 4.**
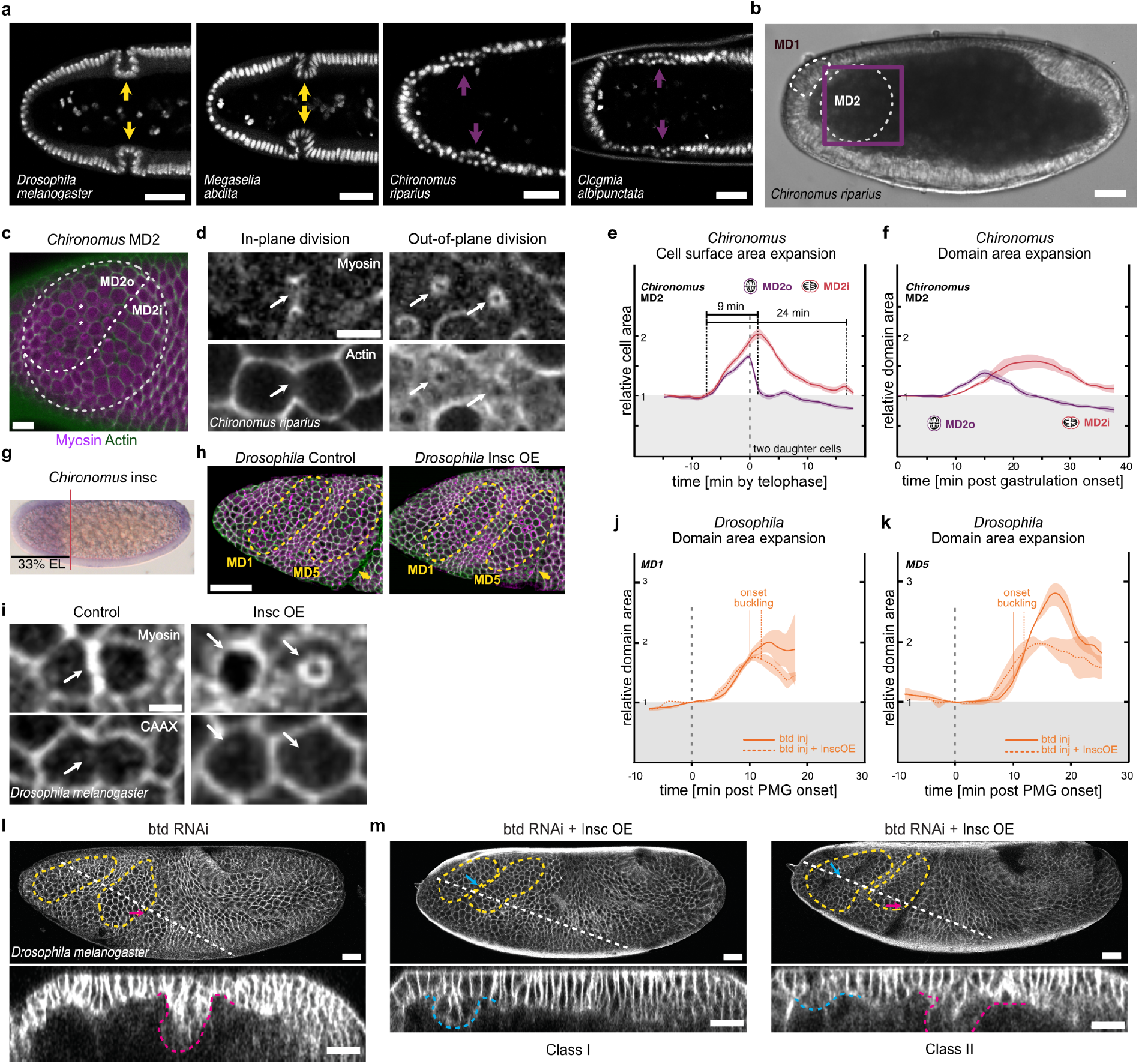
| In flies without a cephalic furrow, out-of-plane cell divisions provide an alternative sink to prevent tissue collision. **a**, Nuclear staining (DRAQ5) in the anterior half of fixed embryos of the 4 representative species with a single row of nuclei in flies with a CF (yellow arrows) and a double row of nuclei (magenta arrows) in flies without CF. n= 13, 13, 24, and 13 embryos respectively for *Drosophila*, *Megaselia*, *Chironomus*, and *Clogmia*. Scale bars: 50 µm for *Drosophila* and *Megaselia*; 25 µm for *Chironomus* and *Clogmia*. **b**, Position of MD1 and MD2 in the *Chironomus* head. Scale bars, 25 µm. **c**, A close-up sum projection of MD2 (purple box in **b**) showing the subdivision of MD2 into MD2o and MD2i by division plane visualized by cell outlines (LifeAct-mCherry) and cytokinetic rings (Sqh-eGFP). Asterisks, the first dividing cells. Scale bars, 25 µm. **d**, Cellular view of the dividing cells in MD2 using Actin and Myosin to visualize cell outlines (LifeAct-mCherry) and cytokinetic rings (Sqh-eGFP). Arrows, cytokinesis with cytokinetic rings perpendicular (in-plane) and parallel (out-of-plane) to the apical surface. Scale bars, 5 µm. **e**, change of individual apical cell surface areas during division for cells dividing out-of-plane (MD2o) and in-plane (MD2i). A value of 1 corresponds to equal area as the mother cell at blastoderm stage, while a value >1 indicates expansion. Following completion of telophase, 2 daughter cells remain on the surface for in-plane divisions (sum plotted), compared to only the more apical daughter cell for out-of-plane divisions. (n=24 out-of-plane and 18 in-plane divisions from 3 embryos) **f**, Change in apical surface area of MD2 subdomains following the onset of gastrulation (n=3 embryos). **g**, *In situ* hybridization showing a broad expression pattern for *Cri-insc* in the head domain (n=3 embryos). For reference, the vertical line indicates the anterior boundary of the *eve1* region at 33%EL similar to Fig. 1k and l. **h**, Lateral surface projection of the *Drosophila* head region in a representative control or InscOE embryo visualized with membrane (3xmScarlet-CaaX, green) and MyoII (Sqh-eGFP, magenta) markers showing MD1 and MD5 (yellow dashed outlines). Yellow arrow, CF. Scale bars, 50 µm. **i**, Cellular view of telophase cells in MD5 showing the cell outlines (3xmScarlet-CaaX) and cytokinetic rings (Sqh-eGFP) in a control (n=5) or InscOE (n=5) embryo. Scale bar, 5 µm. **j, k**, Change in apical surface area of MD1 (**j**) and MD5 (**k**) during mitosis in control (n=3) and InscOE (n=3) embryos that have been injected for *btd* RNAi knockdown. Time is relative to the onset of gastrulation. **l**, **m**, Lateral surface projection (top panels) and z-axis reslice (bottom panels) views of a representative *btd* RNAi embryo (**l**, n=5), or *btd* RNAi embryo with Insc OE (**m**; Class I, n=3; Class II, n=4), visualized with a membrane marker (3xmScarlet-CaaX). Magenta arrows and magenta dashed outlines, head-trunk buckles; cyan arrows and cyan dashed outlines, small buckles near or inside MDs. Scale bars, 30 µm.

To investigate the impact of division orientation on head expansion, we first characterized head mitosis in the *Chironomus* embryo. Live imaging on the lateral side of the embryo reveals that the first cells that divide are located in the anterior-most region of the head, which we designated as MD1 in accordance with the *Drosophila* MD naming convention, where domain number is designated chronologically^13^ (Fig. 4b). Following divisions in MD1, cells in a large lateral domain posterior to MD1 begin to divide, which we designated as MD2 (Fig. 4b). MD2 cells can be seen as beginning to round up as they enter mitosis, ∼7.5 min after the onset of gastrulation, while the first cell that enters telophase does so at ∼13 min post gastrulation onset. These observations indicate that head mitosis occurs soon after the onset of gastrulation and temporally overlaps with trunk expansion, at comparable timing as in *Drosophila* (Fig. 4 Supplement 3a).

We further characterized the cell division plane in the *Chironomus* head. We observed out-of-plane divisions in MD1 (data not shown), but since the MD1 cells are located near the anterior pole where the embryo surface is highly curved and thus challenging to analyze, we focused on the more accessible MD2. Of all the MD2 cells for which the division orientation can be determined unambiguously, about 50% divide out-of-plane (Fig. 4 Supplement 3b). Cells of each division mode occupy a distinct spatial domain, which we termed MD2o and MD2i, respectively, for out-of-plane and in-plane domains (Fig. 4c). MD2o resides in the anterior part of the domain and has an ellipsoid shape; MD2i is posterior and ventral to MD2o and has a crescent shape. The two subdomains undergo division sequentially: the first division occurs in the center of MD2o, from where the subsequent divisions spread out as a concentric wave traveling across the remainder of MD2o, followed by divisions in MD2i (Fig. 4c; Fig. 4 Supplement 3c; Fig. 4 Movie S1). Thus, head mitosis in *Chironomus* differs substantially from that in *Drosophila*: in *Drosophila* all three MDs (MD1, 2, and 5) that are spatial-temporally equivalent to *Chironomus* MD2 display in-plane divisions, while the only head MD that exhibits out-of-plane divisions is the relatively late MD9^13^.

## Out-of-plane divisions show reduced surface expansion and a more rapid release of expansile stress

We hypothesized that in *Chironomus* out-of-plane divisions attenuate the degree of head expansion, thereby avoiding the need to release compressive stress via out-of-plane tissue deformation. To test this possibility, we quantified the temporal dynamics of MD2 surface expansion. As the cell enters mitosis, its apical area increases due to mitotic rounding. Prior to telophase onset, an in-plane dividing on average cell reaches an apical area ∼1.9-fold of the initial area prior to rounding. Following cytokinesis, the two daughter cells reach a combined area of 2-fold, after which they shrink back and occupy a combined surface area identical to that of the mother cell. The total duration of expansion is ∼24 min (Fig. 4e; Fig. 4 Supplement 4a). In contrast, the out-of-plane dividing cell expands only to ∼1.6-fold by telophase for a duration of only ∼9 min. Furthermore, for the daughter cell that remains on the surface following cytokinesis, the occupied surface area decreases rapidly, first to the size of the mother cell and then further down to ∼0.8-fold of the apical area prior to division (Fig. 4e; Fig. 4 Supplement 4b). These results suggest that out-of-plane divisions require smaller surface area, exert a lesser degree of expansile stress to the neighboring cells, and do so over a shorter period of time.

Given that cell divisions within each domain are not fully synchronous (Fig. 4 Supplement 3c), we next measured surface expansion at the tissue level, comparing MD2o with MD2i for ∼30 cells in each domain. We found that MD2o expands for ∼12 min and to a maximum of ∼1.4-fold of the initial area, while MD2i expands for ∼25 min and to a maximum of ∼1.6-fold (Fig. 4f), indicating that the differences observed between in-and out-of-plane divisions in individual cells are conserved at the tissue level. Taken together, our results reveal that the orientation of cell division constitutes a critical parameter that likely controls the accrued compressive stress, suggesting that non-cyclorrhaphan flies use out-of-plane divisions as an alternative mechanism to avoid tissue collision during ectodermal expansion.

## Reorienting head mitosis out-of-plane suppresses CF-region buckling in *Drosophila*

During *Drosophila* gastrulation, spatial-temporally restricted expression of the mitotic spindle anchoring protein Inscuteable (Insc) is required for out-of-plane divisions in MD9^26,27^. We cloned the *Chironomus* ortholog of *insc* and found that it is expressed throughout the entire head region (Fig. 4g). Although we have not been able to assay its functional requirement in division orientation (see Discussion), the expression pattern of *insc* implies that the entire head region of the *Chironomus* embryo is genetically conducive to out-of-plane division.

Taking advantage of the fact that in *Drosophila* Insc is necessary and can be sufficient to instruct out-of-plane division^26^, we overexpressed Insc (Insc^OE^) throughout the head region of the *Drosophila* embryo (see Methods), in an attempt to reorient mitotic spindles in cells that normally exhibit in-plane division to divide out-of-plane. We then asked whether out-of-plane division can replace the reliance on epithelial out-of-plane deformation for dissipation of compressive stress, along with *btd* RNAi to mimic the *Chironomus* head morphogenesis. We focused on MD1 and MD5, and confirmed the altered plane of division through the existence of a circular cytokinetic ring parallel to the embryo surface in most of the cells in these domains (Fig. 4h, i). We then assayed the effect of Insc^OE^ on domain expansion in the *btd* RNAi embryos. For both MD1 and MD5, Insc^OE^ results in a reduced area expansion, i.e. from 2-fold to 1.8-fold in MD1 (Fig. 4j) and from 2.8-fold to 2-fold in MD5 (Fig. 4k). The reduction of area expansion is qualitatively comparable to the differences between in– and out-of-plane divisions in *Chironomus* MD2i and MD2o, suggesting that Insc^OE^ effectively converts the *Drosophila* head into a *Chironomus*-like state, allowing us to ask whether widespread out-of-plane divisions can function as a mechanical sink to release compressive stress.

To test this, we examined the effect of Insc^OE^ and found that although 60% of embryos (Class II) undergo head-trunk buckling comparable to *btd* RNAi alone, 40% of the embryos (Class I) showed near complete loss of the head-trunk buckle (Fig. 4l, m; Fig. 4 Movie S2). Embryos with the Class I phenotype tend to form smaller buckles either in the region between MD1 and MD5 or posterior to MD6, or both. In this context, we know from material science that moderately compressing a thin elastic film residing on a soft compliant substrate results in the formation of short wavelength wrinkles, while further compression beyond a critical threshold pushes the system towards a ‘wrinkle-to-fold transition’, where the deformation becomes localized to a single, deep fold, while all other wrinkles vanish^28,29^. The small buckles that form under Insc^OE^ are reminiscent of the short wavelength wrinkles that form on the elastic film when the compressive load is below the transition threshold, whereas the deep head-trunk buckle may result from wrinkle-to-fold transition. Thus, Insc^OE^ appears to reduce the compressive load to a level near the critical point of fold transition, yielding a divergent phenotype. In support of this, Insc^OE^ decreases CF depth in the wild-type background (Fig. 4 Supplement 5). Thus, Insc^OE^ partially suppresses head-trunk buckling, supporting our hypothesis that orienting mitotic divisions out-of-plane helps dissipate compressive stress to prevent head-trunk collision. In sum, our comparative and functional studies reveal the existence of two programmed morphogenetic solutions to epithelial buckling that can arise due to tissue collision – out-of-plane deformation, in the form of the CF in cyclorrhaphan flies, and out-of-plane division in non-cyclorrhaphan flies.

## Discussion

Our phylogenetic survey provides evidence that the CF, a deep, transient epithelial fold that forms during early *Drosophila* gastrulation, is an evolutionary novelty and a derived character in the monophyletic group of Cyclorrhapha. As such, our data do not support the hypothesis that homologizes the CF with the vertebrate midbrain-hindbrain boundary on the basis of homologous gene expression with deep conservation across phyla^16^. Instead, data in ours and the independent study by Vellutini *et al.*^18^ both support the model that CF formation preempts compressive stress to prevent tissue buckling in the context of a physically confined embryo. As such, the CF executes a mechanical function that provides one possible solution for dipteran gastrulation where the head and trunk tissues expand concurrently. Our data also do not support the hypothesis that the CF functions as an immoble fence to break the symmetry of GBE tissue flow^15^. On the contrary, we find that trunk ectodermal cells are incorporated, and thus ‘flow’, into the CF to increase its depth, indicating that the CF is a sink and a mobile barrier that prevents intermixing of head and trunk tissues. As blocking CF formation causes midline distortion similar to loss of PMG specification, our results suggest that active deformation orchestrated in CF formation, like PMG, buffers stochastic bilateral asymmetry intrinsic to GBE^30^. Furthermore, our phylogenetic mapping reveals an abrupt and near concurrent emergence of the two morphogenetic traits: predominantly in-plane division for the early mitotic cells in the head and active out-of-plane deformation at the head-trunk boundary (the CF). Notably, species that represent the most basal branches of Cyclorrhapha, such as Megaselia, contain both of these traits, raising the intriguing question regarding which of them evolved first.

If in-plane division arose first, the increased spatial demand and the prolonged surface expansion associated with in-plane division would likely have exerted stronger compressive stress on the head-trunk boundary, increasing the likelihood that it would buckle. Given that buckling and normal GBE are both intrinsically stochastic and can increase the probability of bilateral asymmetry^30^ (Fig. 3), it is likely that organisms with in-plane head divisions could have only arisen and survived initially under conditions that were more favorable to stress dissipation, e.g. slower developmental rate or cooler temperature, tempering the likelihood of axial distortion. Under such conditions, the presence of genetically programmed out-of-plane deformation, such as CF formation, would have likely increased developmental robustness and allowed adaptation to a wider range of developmental and environmental conditions. This hypothesis predicts that reorienting head division from primarily out-of-plane to in-plane can cause ectodermal buckling in *Chironomus*. Unfortunately, testing this hypothesis has been challenging, as our preliminary attempt at reorienting the division plane in *Chironomus* via downregulation of Insc expression has not been successful, suggesting that early anterior expression of *insc* is just one of several redundant mechanisms to orient the division plane in *Chironomus*. Additional cellular components and mechanical conditions may include high surface tension in the mitotic cortex and dedicated cues for anchorage of mitotic spindle at the lateral cortex^31–34^. The evolutionary transition to in-plane division would thus require more than a mere restriction of *insc* expression. Indeed, the presence of both out– and in-plane dividers within the *Chironomus* head MDs despite the presence of Insc suggests potential volatility of division orientation in the last common ancestor of cyclorrhaphan and non-cyclorrhaphan flies.

If the CF evolved first, arising prior to the conversion of predominant in-plane divisions in the head, compressive stresses in the head may have been initially low or absent, and the CF may have arisen for a different, currently unknown function. In this scenario, active folding of a CF at the head-trunk boundary would have pulled on the neighboring tissue, making it energetically more favorable for cells in the head region to divide in-plane, potentially paving the way for an eventual evolutionary transition to obligatory in-plane division, e.g. by the loss of early *insc* expression in the head and the gain of additional cellular and mechanical conditions that can promote in-plane division. This scenario posits that the CF was co-opted secondarily to function as a mechanical sink, following the evolution of in-plane divisions in the head.

Support for this could come from identification of additional functions of the CF during development. Previous studies provided some hints. For example, one hypothesis proposes that the CF serves as a temporary storage of cells that will subsequently contribute to future head development^35^, while lineage tracing experiments suggest that the MD2 cells that form a part of the CF give rise to the floor of the pharynx and associated macrophages or muscle cells^36^. It would thus be worthwhile employing methods established in our work and the independent study by Vellutini *et al*.^18^ to further explore the possibility that the CF carries out additional functions in *Drosophila*, *Megaselia*, and potentially other species representing the basal branches of Cyclorrhapha.

CF formation in *Drosophila* has been previously shown to represent a morphogenetic process whose spatial precision depends on not only the robustness of genetic patterning, but also mechanical self-organization^9^. Our surprising findings of the CF as an evolutionary novelty, and its co-evolution with the reorientation of the plane of mitotic division in a neighboring tissue, may provide a unique window to peek into the role of mechanics in the evolution of morphogenetic processes. In particular, our work raises the possibility that inter-tissue mechanical conflict, or mechanical constraint in general, constitutes a mechanism of positive feedback^37^ that could drive rapid evolutionary transitions and the seemingly abrupt emergence of morphogenetic traits.

## Figure supplements

**Fig. 1 Supplement 1.**
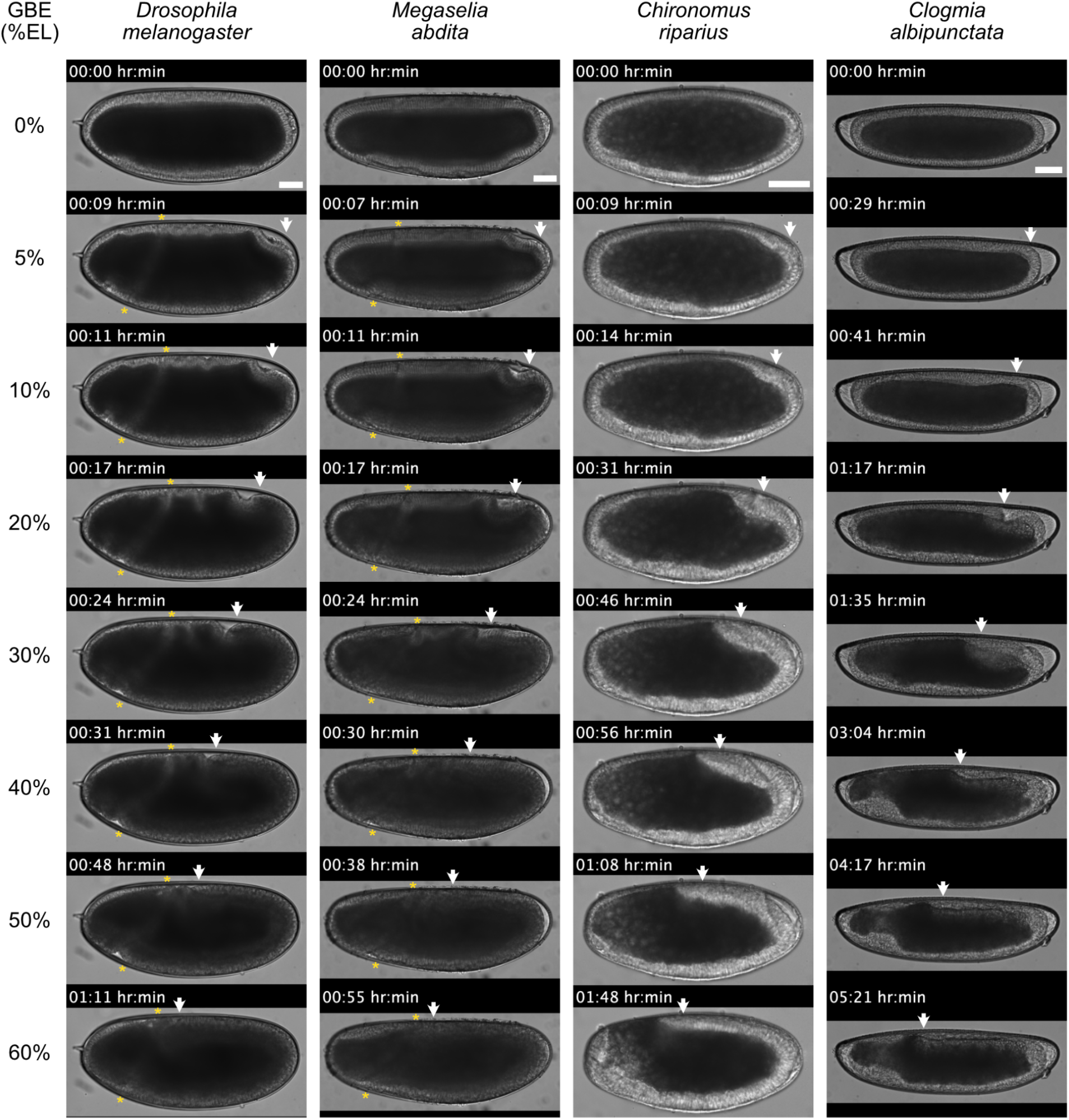
| Progression of GBE is comparable in all species, while only *Drosophila* and *Megaselia* show a CF. Figure shows montages of representative DIC recordings of developing embryos from our four species of interest. T_0_ is defined as the initiation of gastrulation. Asterisks mark/track the CF in *Drosophila* and *Megaselia*, while the arrow tracks the posterior end of the ectoderm in all species. n= 35, 24, 27, and 42 embryos respectively for *Drosophila*, *Megaselia*, *Chironomus*, and *Clogmia*. Scale bars 50 µm.

**Fig. 1 Supplement 2.**
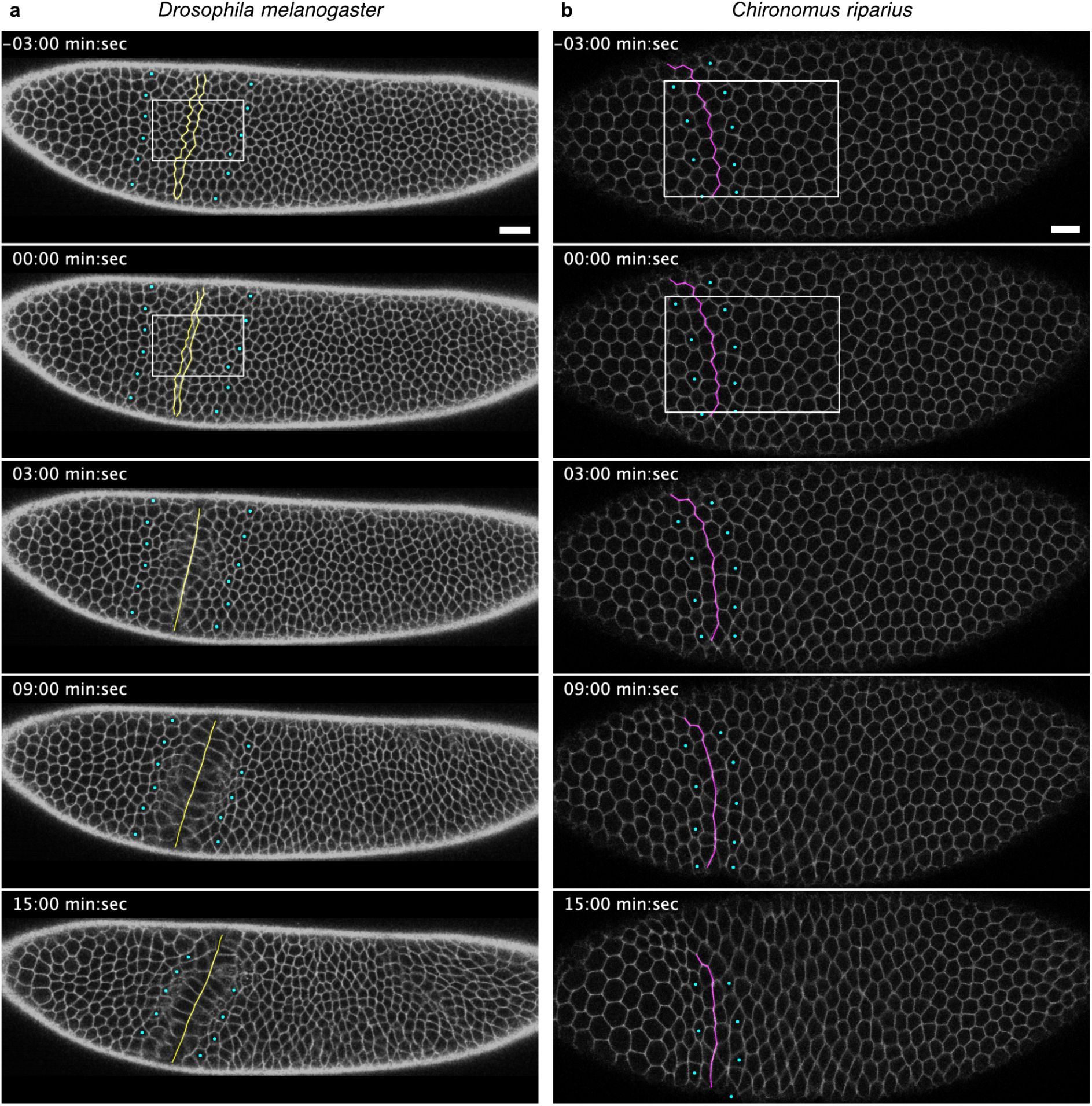
| *Chironomus* does not form a CF at the head-trunk boundary. **a** and **b**, Montage of snapshots from a single z-slice over time, from live imaging *Drosophila* (**a**, n=3) and *Chironomus* (**b**, n=3) embryos, expressing cell membrane markers. In the examples shown, the regions correspond to ∼10 to 75% EL for *Drosophila* and ∼15 to 85% EL for *Chironomus*. T_0_ is defined as initiation of gastrulation. The jagged colored lines mark the cell boundaries between head and trunk ectoderm. The boundaries are inferred from the appearance of CF in *Drosophila* embryo, while in case of *Chironomus* embryo the differences in cell shape changes dictate the placement of the boundary. Cyan dots track cells that can be followed from the first frame and end-up close to the head-trunk boundary in the last frame. While the straightening of the jagged lines is comparable in both species, the subsequent cell shape changes prior to formation of the CF and the tissue flow towards the head-trunk boundary are absent in *Chironomus*. Boxes indicate the regions shown in Fig. 1g and h. Scale bar, 20 µm *Drosophila*, 10 µm *Chironomus*.

**Fig. 1 Supplement 3.**
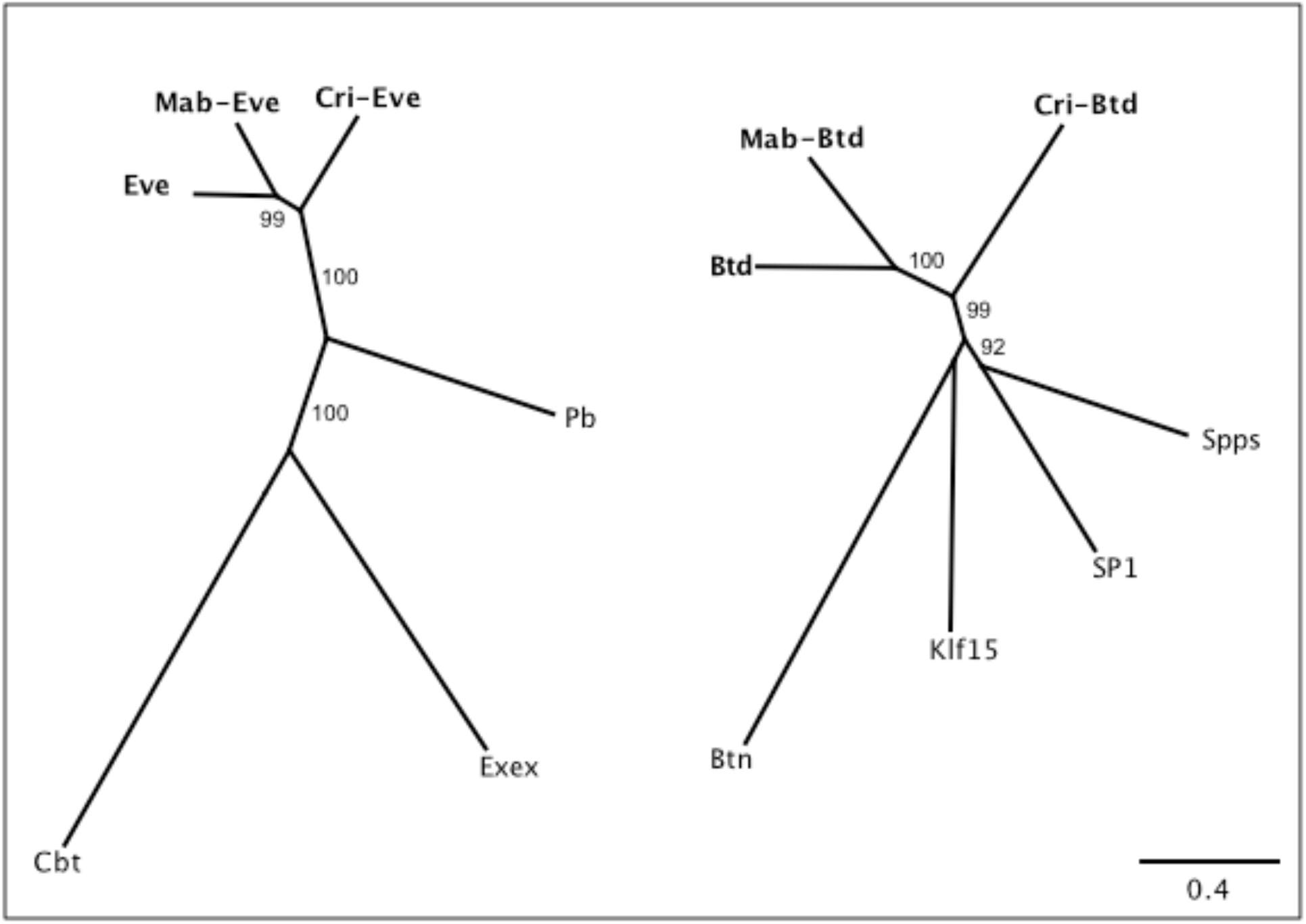
| Protein tree of predicted orthologs of *eve* and *btd* in *Megaselia* and *Chironomus*. Phylogenetic distances of Even-skipped (Eve) and Buttonhead (Btd), its predicted orthologs in *Megaselia* and *Chironomus*, as well as the three most closely related proteins in *Drosophila* were calculated in Geneious using Jukes-Cantor as the genetic distance model. Numbers refer to reliability values in percent, with values shown only above 80. RefSeq protein sequences from NCBI for *Drosophila melanogaster* are Btd (Buttonhead, NP_511100), Btn (Buttonless, NP_732768), Cbt (Cabut, NP_722636), Eve (Even-skipped, NP_523670), Exex (Extra-extra, NP_648164), Klf15 (Kruppel-like factor 15, NP_572185), Pb (Proboscipedia, NP_476669), SP1 (SP1, NP_727360), and Spps (SP1 like factor, NP_651232). Scale bar is the number of changes per site.

**Fig. 1 Supplement 4.**
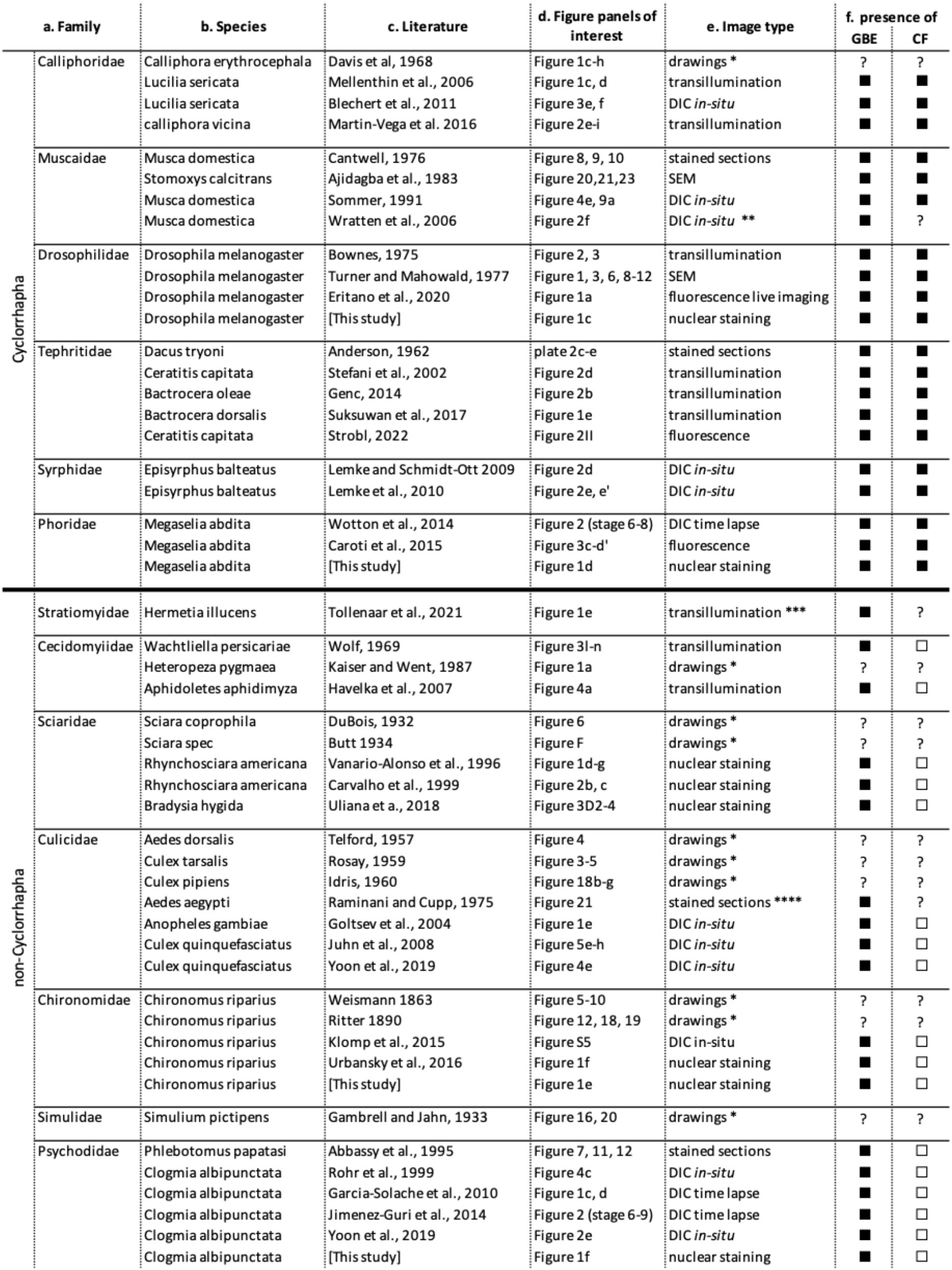
| Literature survey for the presence or absence of CF in diptera. This supplementary table compiles a list of literature references where evidence for the presence or absence of CF and GBE in a broad range of species encompassing the whole order of Diptera could be found. **a**, Fly families in the order of Diptera for which references could be found. **b** and **c**, Names of the species and the corresponding references. **d**, Reference figure panels with images of embryos at various stages of early gastrulation that were used to interpret the presence or absence of GBE and CF. **e**, Description of image datatype in **d.** SEM, scanning electron micrographs; DIC, Differential Interference Contrast imaging; *in-situ*, whole mount micrographs of embryos from RNA *in-situ* hybridization experiments. **f**, Interpretation of the presence or absence of GBE and CF based on images in **d**. Black boxes, presence; hollow boxes, absence; question marks, unclear (see comments below). *, Descriptive drawings are a classic resource for the depiction of embryonic development. However, we have refrained from using these as definitive evidence for the presence or absence of GBE and/or CF in a given family, due to the potential for subjective interpretations^58^. **, unclear evidence for CF due to absence of images during early gastrulation. Note a late furrow is visible in the image, though the stage is too late to fulfill the definition of CF. ***, unclear evidence for CF due to absence of images during early gastrulation. Although no furrow is visible during late gastrulation, presence of an early transient furrow can not be excluded. ****, unclear evidence for CF due to absence of images from the anterior region of the embryo.

---

**Fig. 2 Supplement 1.**
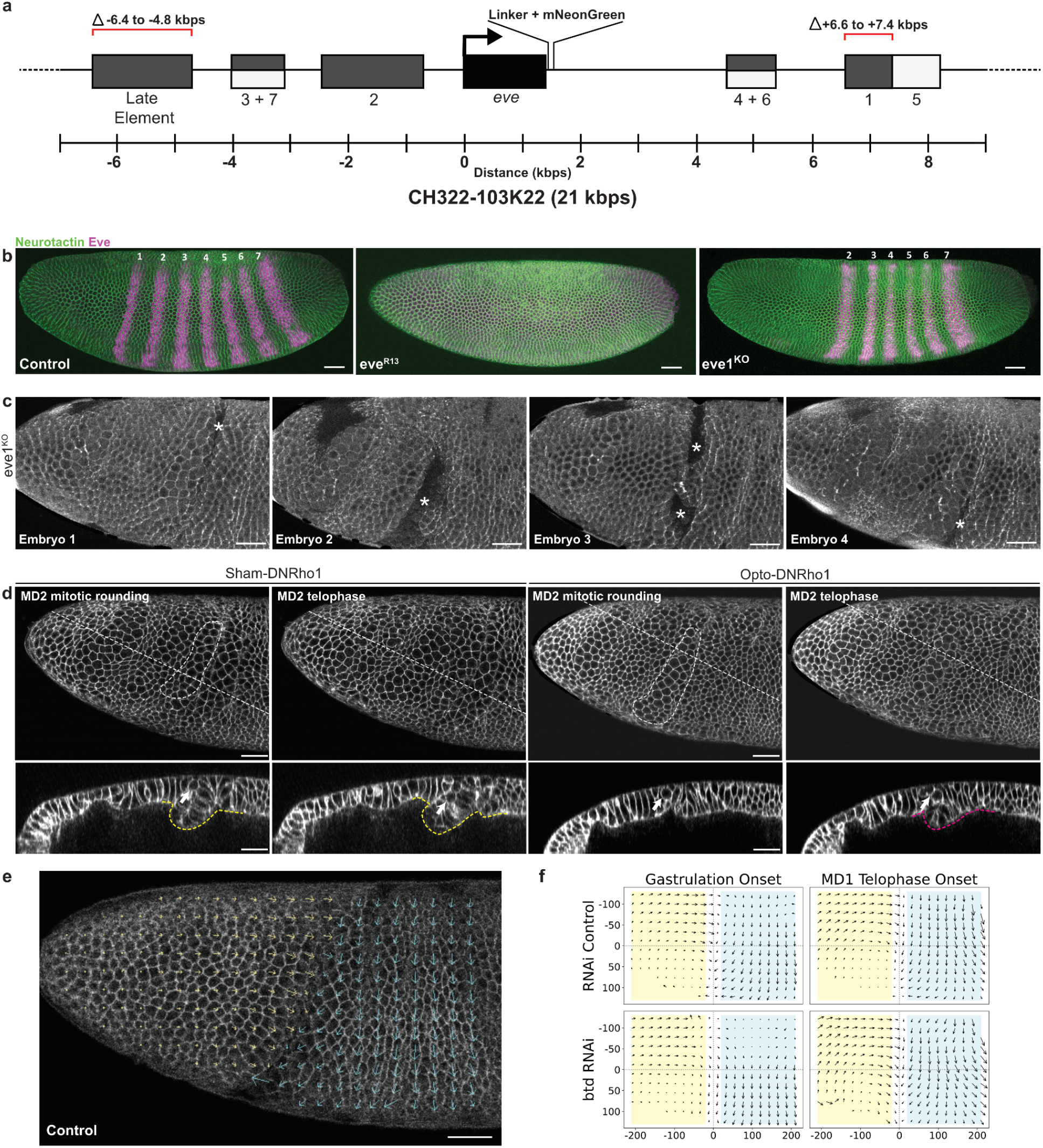
| Phenotypic analysis following genetic or optogenetic ablation of the CF. **a**, Schematic representation of the *eve^CH322–103K22^-mNeonGreenΔst1ΔLE* construct. *eve*, transcriptional region of the *eve* gene; numerics (1, 2, 3+7, 4+6 and 5) and Late element, enhancer regions; scale, genomic coordinates relative to the transcriptional start of *eve*. **b**, Lateral surface projection of membrane (labeled with anti-Neurotactin, green) and Eve (magenta) immunofluorescence in a representative control (n=5), *eve^R13^* (n=10) or *eve1^KO^* (n=14) embryo imaged on the lateral side. Numerics, the Eve stripes. **c**, Lateral surface projection of four representative *eve1^KO^* embryos (n=4) expressing MyoII-mKate2 showing variable D-V positions of the initial head-trunk buckles (white asterisks). **d**, Lateral surface projection of a representative sham control (Sham-DNRho1) or a photoactivated (Opto-DNRho1) embryo expressing the Opto-DNRho1 system, visualized with a membrane marker (3xmScarlet-CaaX) showing the anterior half of the embryo that includes the head-trunk boundary (top rows) and a z-axis reslice (bottom rows) at two time points. White dashed outlines and white arrows, rounding or dividing MD2 cells; yellow outlines, CFs; magenta outline, head-trunk buckle. Note in the sham control MD2 rounding occurs when the out-of-plane deformation of the CF was already present, while the onset of head-trunk buckle coincided with MD2 rounding when the CF was eliminated via optogenetic inhibition of DNRho1. **e**, Representative image of a control embryo superimposed with tissue flow field visualized with PIV. Yellow and cyan arrows are vector fields of the head and trunk regions, respectively. Scale bar, 30 µm. **f**, Tissue flow fields visualized with PIV in RNAi control (n=4) and *btd* RNAi (n=5) embryos at the onset of gastrulation and MD1 telophase. Yellow shaded rectangles, head region; blue shaded rectangles, trunk regions; x-origin, head-trunk boundary; y-origin, lateral midline. Unit, arbitrary unit.

**Fig. 2 Supplement 2.**
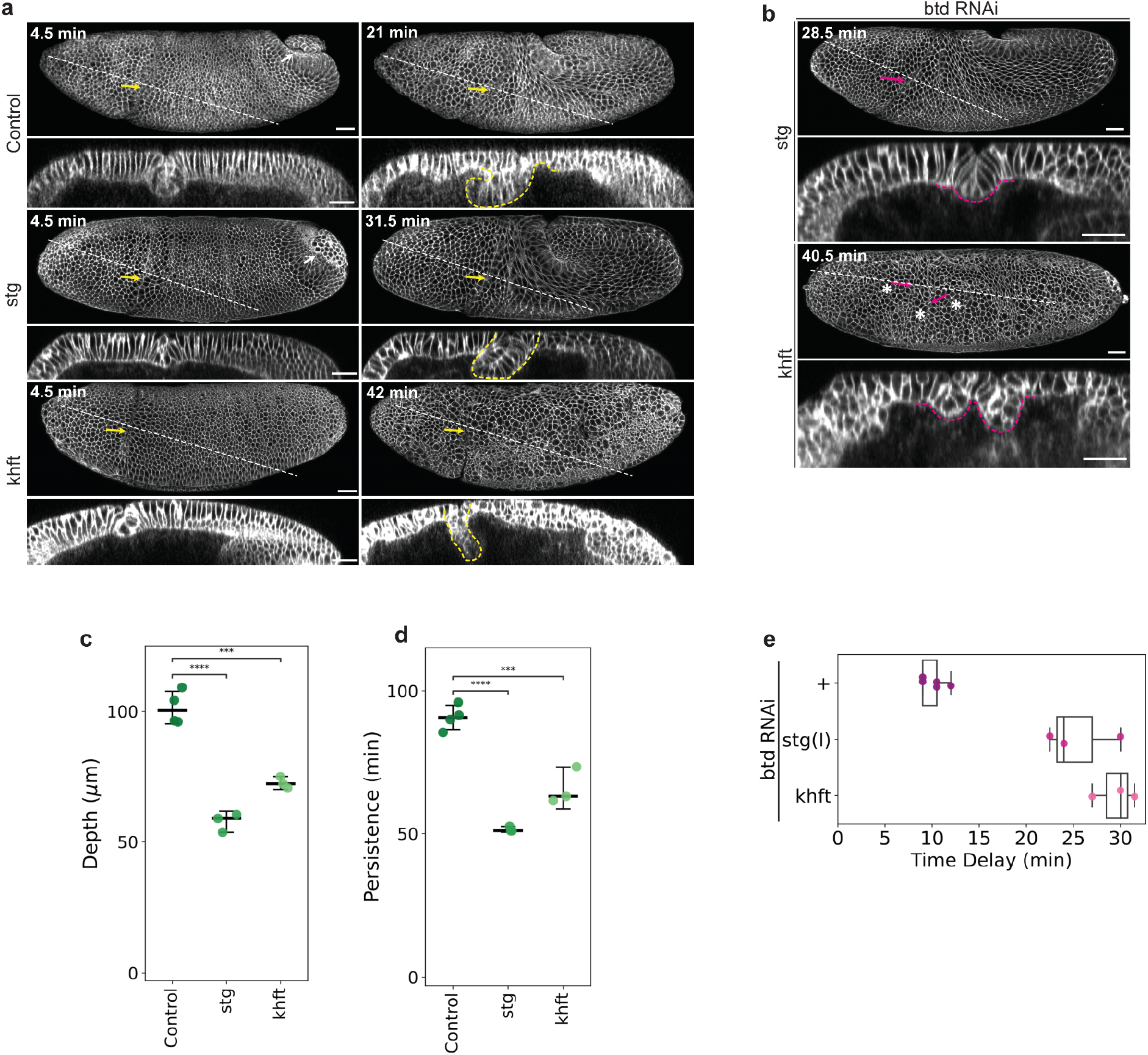
| The effects of head mitosis and trunk convergent-extension on the CF and head-trunk buckling when the CF is lost. **a**, Time-lapse series of a representative control (*mat-tub-3xmScarlet-CaaX/+* with water injection, n=4), *stg* mutant (n=3), or *khft* (n=5) mutant embryo, visualized with 3xmScarlet-CaaX showing a lateral surface projection (tow rows) and a z-axis reslice (bottom rows). White arrows, PMG; yellow arrows and yellow dashed outlines, CFs. Time is relative to the onset of gastrulation. Scale bars, 30 µm. **b**, Lateral surface projection (top panels) and z-axis reslice (bottom panels) of 3xmScarlet-CaaX showing late-stage buckling in *stg* (n=3) or *khft* (n=3) mutants (average buckle depth 40.31± 5.31 µm). Magenta arrows and magenta dashed outlines, head-trunk buckles; asterisks, late MDs (likely MD5, 6, and 11, based on their location, though the timing appears abnormal). Time is relative to the onset of gastrulation. Scale bars, 30 µm. **c**, **d**, Maximum depths (**c**) or durations (**d**) of the CF plotted with median and error bars indicating 95% confidence interval. One-way ANOVA Tukey post-hoc test; ****, p < 0.0001; ***, p < 0.001. Sample size: control=4, *stg*=3, *khft*=3. **e**, Timing of the onset of late buckling in the *stg*(I) and *khft* embryos that have been injected with *btd* RNAi, as compared to *btd* RNAi alone. Time is relative to the onset of gastrulation. Sample size: Control (*btd* RNAi alone)=5, *stg*(I)=3, *khft*=3.

**Fig. 3 Supplement 1.**
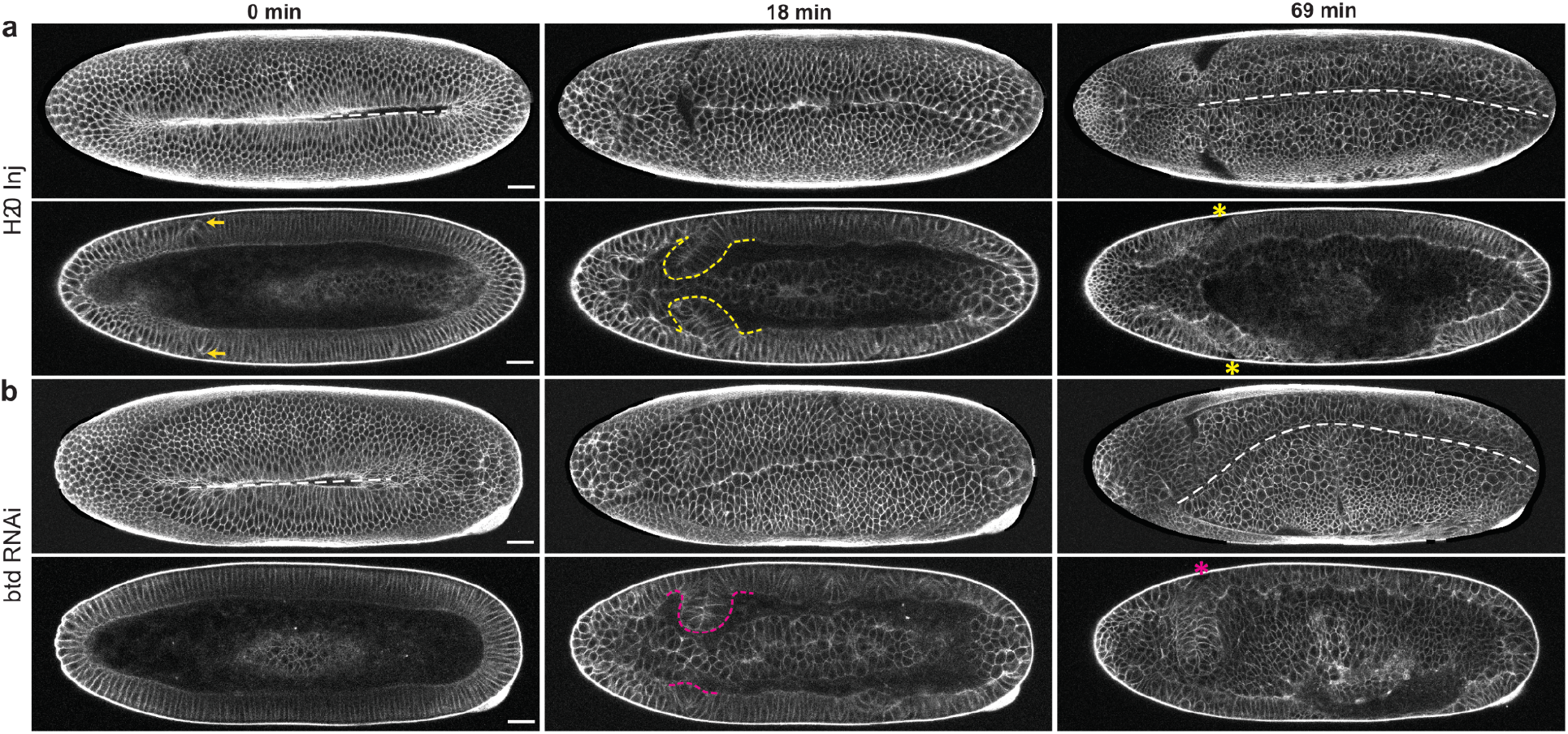
| Elimination of the CF via *btd* RNAi also causes midline distortion. **a**, **b**, Time-lapse series of a representative water injection control (**a**, H_2_O inj, n=5) or *btd* RNAi (**b**, n=5) embryo, visualized with a 3xmScarlet-CaaX showing a ventral surface projection (top rows) and a single coronal section (bottom rows). White dashed lines, ventral midlines; yellow arrows and yellow dashed outlines, CF; yellow asterisks, bilaterally symmetric CFs; magenta arrows and magenta dashed outlines, head-trunk buckles; magenta asterisks, bilaterally asymmetric buckles. Time is relative to the onset of gastrulation. Scale bars, 30 µm.

**Fig. 4 Supplement 1.**
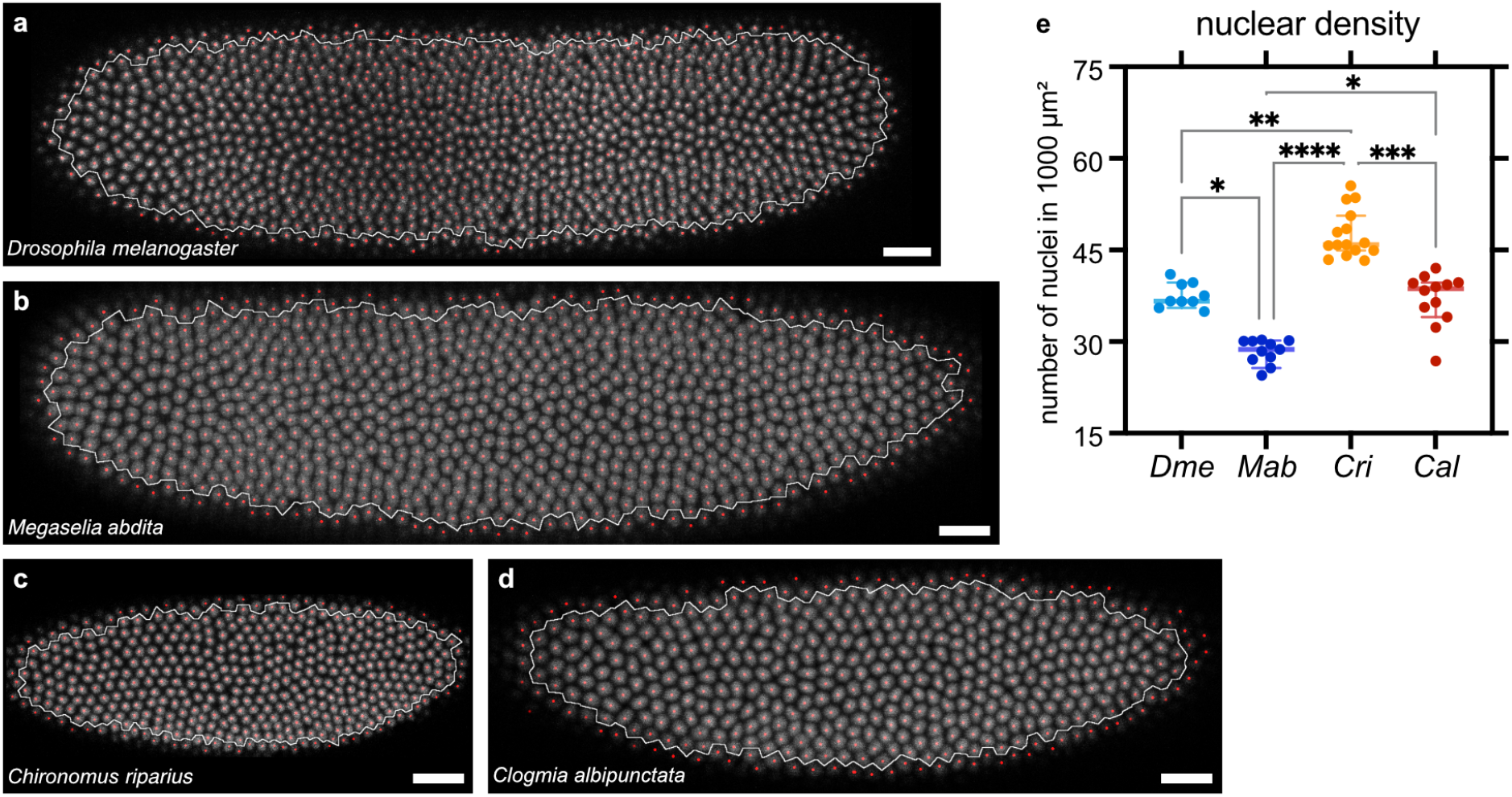
| Blastoderm cell density is comparable across species. **a**-**d**, Shown are representative images from fixed embryos of our species of interest, stained with DRAQ5 (see methods), and segmented nuclei; nuclei are shown in gray-scale, with semi-automatically generated red dots as tags for individual nuclei. The distribution of the dots is used to compute the white boundary. The nuclear density is estimated as the number of red dots divided by the area of the region bounded by the white boundary. Scale bars, 20 µm. **e**, Plot shows the distribution of nuclear density, with each dot representing data from one embryo, bold horizontal line indicating median, and whiskers showing 95% confidence intervals. Embryos were pooled from at least 2 fixation and staining rounds. n= 9, 11, 15, and 12 embryos respectively for *Drosophila*, *Megaselia*, *Chironomus*, and *Clogmia*. The trends in the changes in nuclear density in these species do not match with presence or absence of a CF. *, p<0.05; **, p<0.01; ***, p<0.001; ****, p<0.0001; non-significant differences are omitted.

**Fig. 4 Supplement 2.**
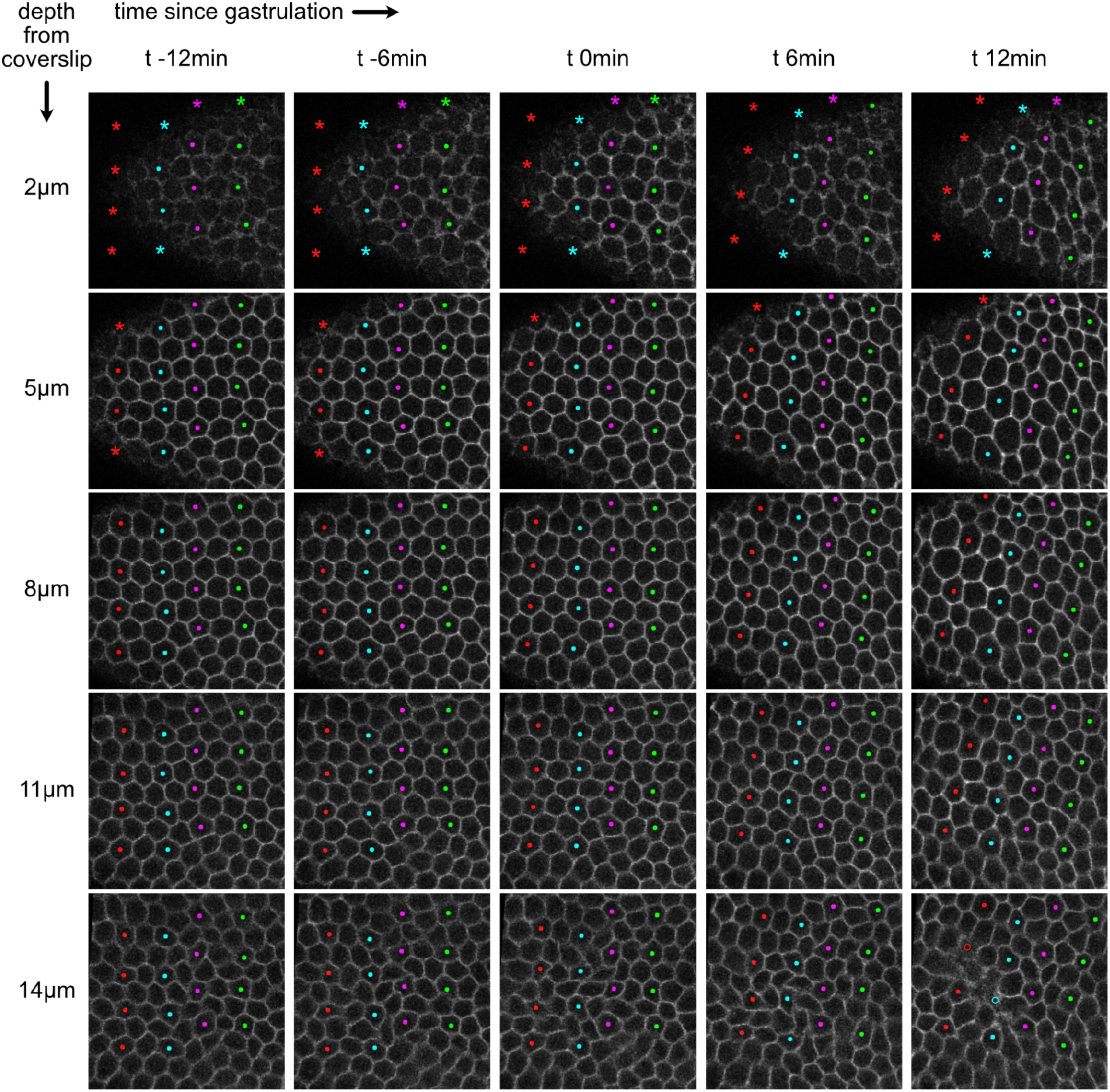
| Distribution of nuclei in 2 layers can not be explained by delamination or pseudostratification. Time lapse recordings of cell behavior in the head of early *Chironomus* embryos to test whether the two layers of nuclei observed in the head region of fixed *Chironomus* and *Clogmia* embryos (Fig. 4a) might stem from pseudostratification or cell extrusion. In either of the scenarios, a fraction of nuclei are known to be basally located, and we would expect to find wedge or cone shaped cells ^38,39^. Accordingly, we reject either alternative explanations as we observe cells to be columnar till they begin mitotic rounding. Montages show different z-stacks arranged along columns, while different time points arranged along rows, in the head region of a representative *Chironomus* embryo (n=3). Each panel is 50×50 µm. In total 16 cells are marked in all of the panels using colored dots. Some cells could not be seen in the z-stacks near the surface due to the curvature of the embryo, and in such cases asterisks approximate the cell position. Some cells could not be seen in the z-stacks in basal regions due to mitotic rounding as they retract from the basal side, and in such cases hollow markers approximate the cell position deep inside the embryo.

**Fig. 4 Supplement 3.**
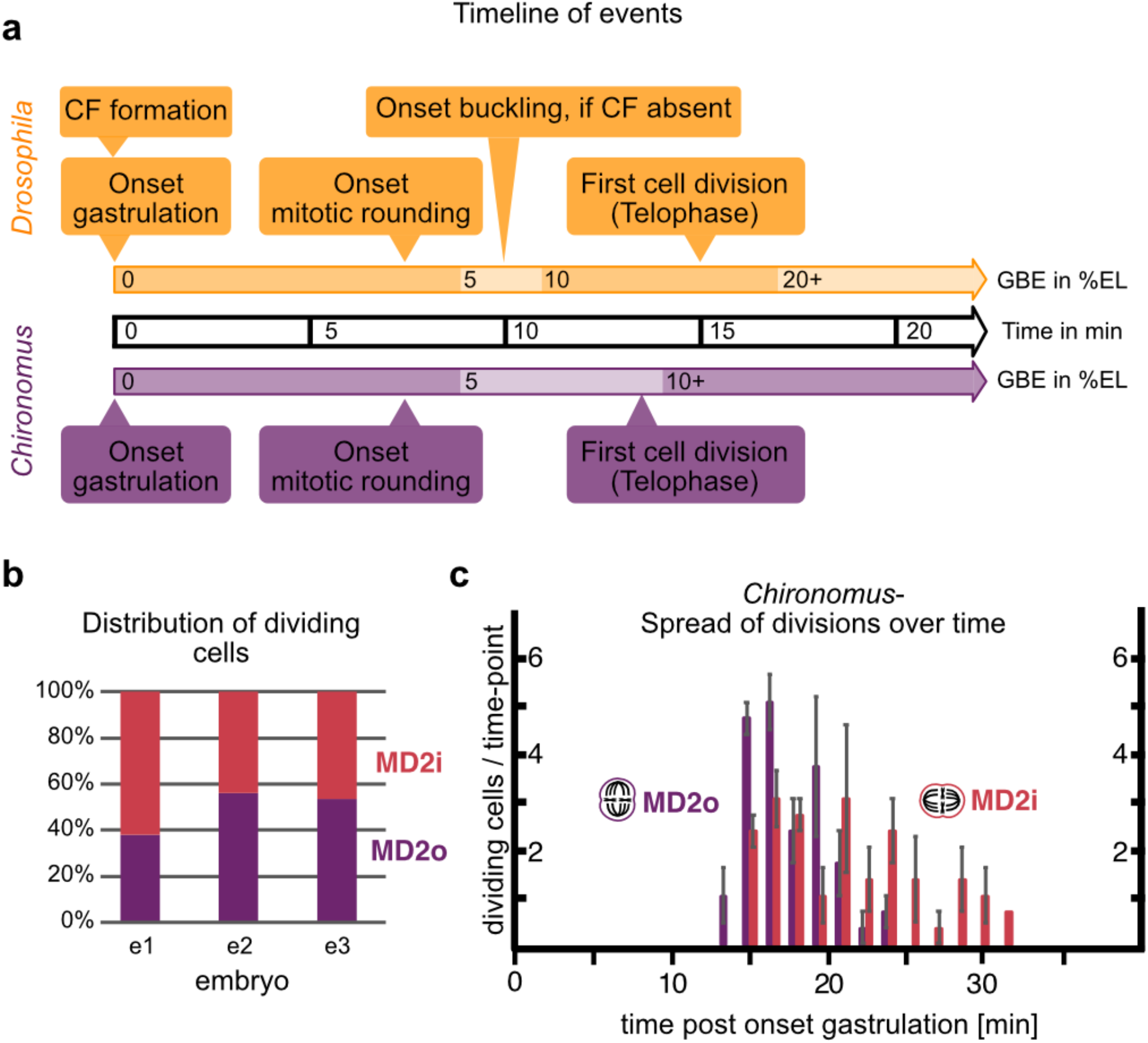
| Division event timeline in *Drosophila* and *Chironomus* and spread of divisions over time in *Chironomus* MD2. **a**, Timeline of division events (yellow-*Drosophila*, magenta-*Chironomus*) in absolute time (in min) post gastrulation onset, defined as the onset of PMG in *Drosophila* and the onset of ventral-ward tissue flow in *Chironomus*. The onset of mitotic rounding was determined by the first cell displaying rounding in the domains analyzed (MD1 in *Drosophila*, MD2o in *Chironomus*). Complementary, the degree of GBE by % egg length (EL) in this timeframe is given for each fly. **b**, Distribution of cells dividing out-of-plane (magenta/MD2o) and in-plane (red/MD2o) in the 3 embryos analyzed.**c**, Spread of divisions over time in *Chironomus* MD2o and MD2i in time post onset of gastrulation. Bars represent the number of cells entering telophase within a 90s window. n=3 embryos.

**Fig. 4 Supplement 4.**
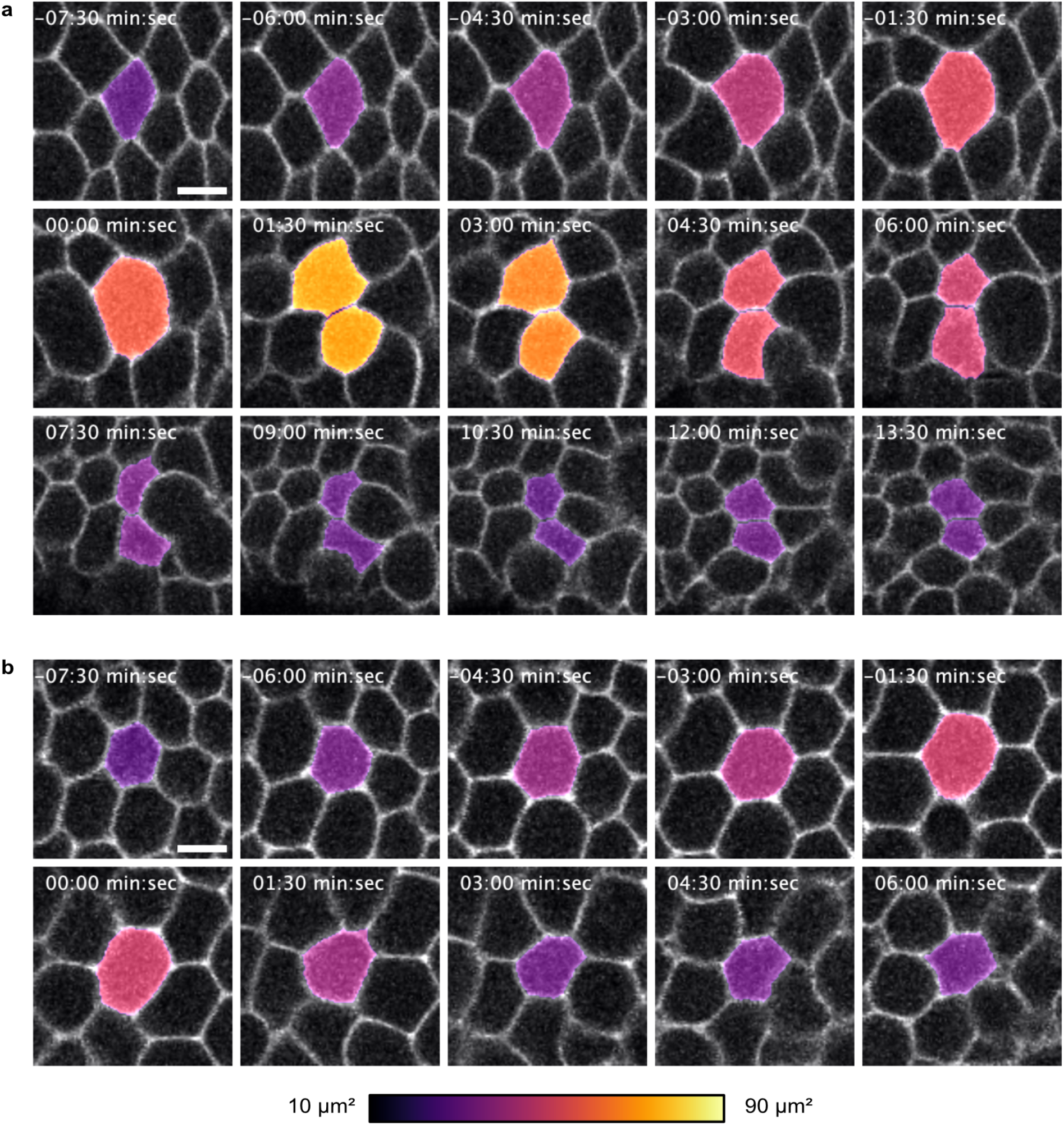
| Time course of an out-of-plane vs. in-plane division, with associated changes in apical cell area in *Chironomus* embryos. **a** and **b**, Time course of an in-plane division (**a**) and an out-of-plane division (**b**) with T_0_ defined the same way as that in Fig. 4e, i.e., the onset of telophase. Cells are chosen from a representative embryo, **a** from MD2i and **b** from MD2o. The LUT bar at the bottom indicates the color-code used for cell apical area. Scale bars 5 µm.

**Fig. 4 Supplement 5.**
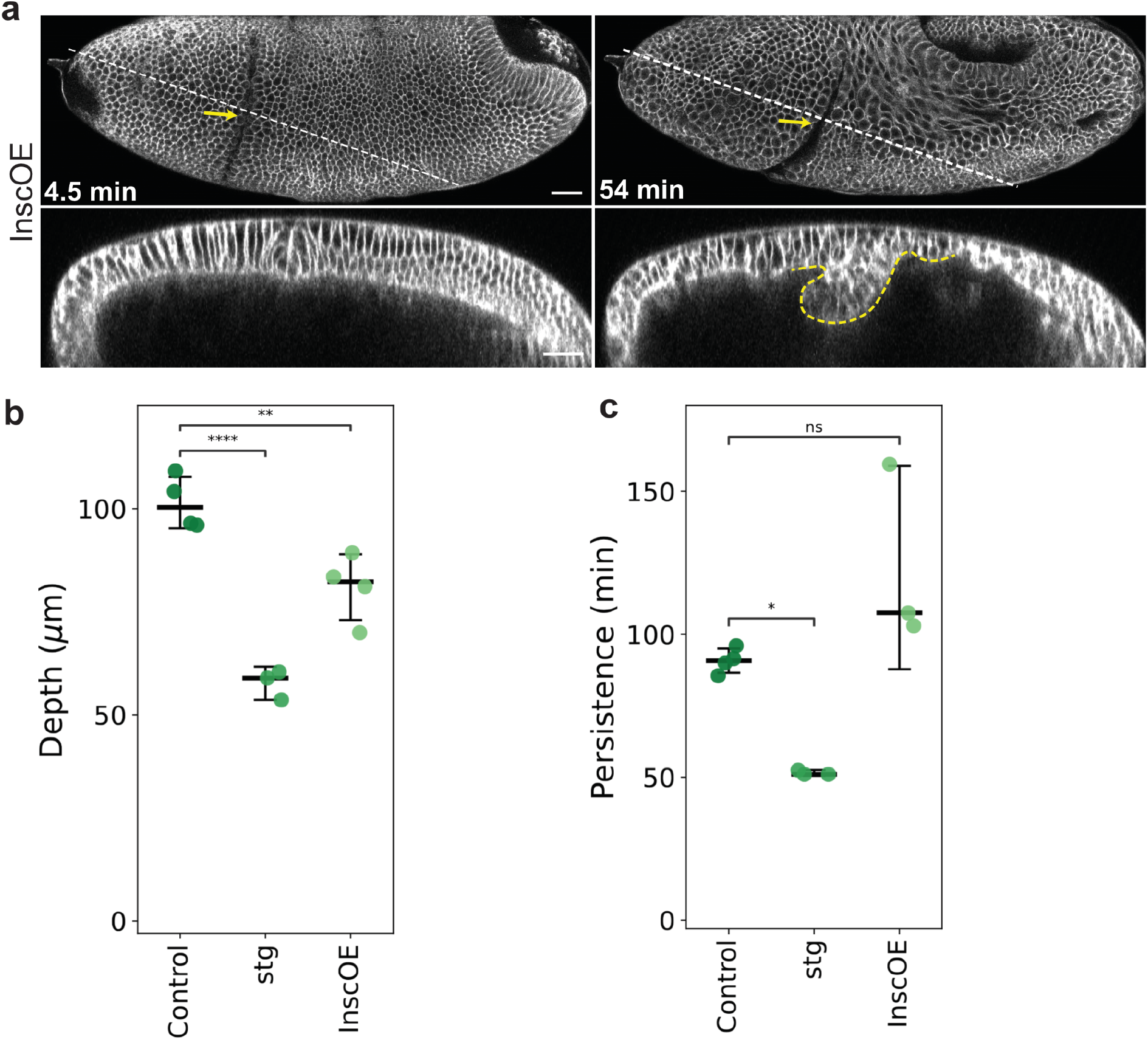
| The effect of Insc overexpression on CF formation. **a**, Time-lapse series of a representative embryo of Insc overexpression in the head region (InscOE, n=5), visualized with 3xmScarlet-CaaX showing a lateral surface projection (top rows) and a z-axis reslice (bottom rows). Yellow arrows and yellow dashed outlines, CFs. Time is relative to the onset of gastrulation. The CF reaches the maximal depth at 54 min. Scale bars, 30 µm. **b**, **c**, Maximum depths (Sample size: control=4, *stg*=3, InscOE=4) (**b**) or durations (Sample size: control=4, *stg*=3, InscOE=3) (**c**) of the CF plotted with median and error bars indicating 95% confidence interval. One-way ANOVA Tukey post-hoc test; ****, p < 0.0001; **, p < 0.01; *, p < 0.1. Control and *stg* measurements were replicated from Fig. 2 Supplement 2c, d.

## Figure legends [movies]

**Figure 1 Movie S1 | *Chironomus* does not form a CF at the head-trunk boundary**

Lateral view from a single z-slice of a developing *Drosophila* (top panel) and *Chironomus* (bottom panel) embryos, expressing cell membrane markers. The regions correspond to ∼10 to 75% EL for *Drosophila* and ∼15 to 85% EL for *Chironomus*. The colored lines mark the cell boundaries between head and trunk ectoderm. The boundaries are inferred from the appearance of CF in *Drosophila* embryo, while in case of *Chironomus* embryo the differences in cell shape changes dictate the placement of the boundary. Cyan dots track cells that can be followed from the first frame and end-up close to the head-trunk boundary in the last frame. While the straightening of the jagged lines is comparable in both species, the subsequent cell shape changes prior to formation of the CF and the tissue flow towards the head-trunk boundary are absent in *Chironomus*. Fig. 1g and h show time points –3 min and 0 min in the boxed region. We are presenting extra timepoints in the *Chironomus* embryo to compensate for slower development. Scale bar, 20 µm *Drosophila*, 10 µm *Chironomus*.

**Figure 2 Movie S1 | *eve1^KO^* lacks MyoII planar polarization in the pre-CF domain**

Lateral surface projection of the head region (top panels) and z-axis reslice view (bottom panels) of control and *eve1*^KO^ embryos expressing MyoII-mKate2. White rectangle outlines MyoII planar polarization preceding CF initiation, while no clear MyoII polarization was observed in *eve1*^KO^. Yellow arrow, CF initiation; magenta arrow, onset of buckling. Time is relative to the onset of gastrulation. Scale bar, 30 µm.

**Figure 2 Movie S2 | Genetic ablation of the CF leads to buckling in the head-trunk boundary**

Lateral surface projection of control, *eve1^KO^*, *btd^AX^* and *eve^R13^* embryos expressing Gap43-mCherry. White arrows, PMG; yellow arrows, CFs; magenta arrows, head-trunk buckles. Time is relative to the onset of gastrulation. Scale bars, 30 µm.

**Figure 2 Movie S3 | Blocking CF formation via optogenetic inactivation of MyoII leads to head-trunk buckling**

Lateral surface projection of the head region and z-axis reslice view of embryos expressing the opto-DNRho1 system, visualized with 3xmScarlet-CaaX, each of which was illuminated with 0% (Sham-DNRho1) or 0.1% (Opto-DNRho1) 405 nm laser light within the blue rectangle. Yellow arrows, CFs; magenta arrows, buckles. Time is relative to the onset of gastrulation. Scale bars, 30 µm.

**Figure 2 Movie S4 | CF initiation occurs normally in *stg* and *khft* embryos, while buckling in the *btd* RNAi embryo is suppressed when mitosis or GBE is abrogated (via *stg* or *khft* mutant)**

Lateral surface projection of control, *stg* and *khft* embryos (left column) and *btd* RNAi, *btd* RNAi+*stg* and *btd* RNAi+*khft* embryos (right column), all visualized with 3xmScarlet-CaaX. White arrows, PMG; yellow arrows, CFs; magenta arrows, buckles. Time is relative to the onset of gastrulation. Scale bars, 30 µm.

**Figure 2 Movie S5 | Reslice view of *btd* RNAi embryos reveals head-trunk buckling is suppressed when mitosis or GBE is abrogated (via *stg* or *khft* mutant)**

Z-axis reslice view of *btd* RNAi, *btd* RNAi+*stg* and *btd* RNAi+*khft* embryos, visualized with 3xmScarlet-CaaX. Time is relative to the onset of gastrulation. Scale bars, 30 µm.

**Figure 3 Movie S1 | Bilateral elimination of the CF via optogenetic inhibition of MyoII causes ventral midline distortion**

Ventral surface projection of embryos expressing the opto-DNRho1 system, visualized with 3xmScarlet-CaaX, each of which was illuminated with 0% (Sham-DNRho1) or 0.1% (Opto-DNRho1) 405 nm laser light within the blue rectangles. White lines, ventral midline. Time is relative to the onset of gastrulation. Scale bars, 30 µm.

**Figure 3 Movie S2 | Genetic ablation of the CF via *btd* mutant also causes ventral midline distortion**

Ventral surface projection of control and *btd*^AX^ embryos expressing 3xmScarlet-CaaX. White lines, ventral midline. Time is relative to the onset of gastrulation. Scale bars, 30 µm.

**Figure 4 Movie S1 | Temporal dynamics of divisions in the *Chironomus* MD2o and MD2i**

Lateral view of *Chironomus riparius* head in embryos injected with Gap43-eGFP mRNA. Dividing cells are marked 5 timepoints before their division and 3 timepoints after division. Out-of-plane divisions (MD2o) are colored magenta to cyan and in-plane divisions (MD2i) are red to yellow. Scale bar is 10 µm.

**Figure 4 Movie S2 | Mitotic rounding and cytokinetic ring positioning during cell division in *Chironomus riparius* in and out-of-plane divisions**

Cellular view of *Chironomus riparius* MD2 cells dividing in different planes. Dividing cells are indicated with a cell outline (gray) based on sum projections of actin (LifeAct-mCherry/green) with cytokinetic ring visualized by Sqh-eGFP (magenta). Representative cells were chosen. Scale bars 5 µm.

**Figure 4 Movie S3 | InscOE reorients the division plane in MD1 and MD5 to out-of-plane division**

Lateral surface projection of the head region of control and *InscOE* embryos expressing Sqh-eGFP (magenta) and 3xmScarlet-CaaX (green). Time is relative to the onset of gastrulation. Scale bars, 30 µm.

**Figure 4 Movie S4 | Two classes of phenotypes in InscOE + *btd* RNAi**

Lateral surface projection (top panels) and z-axis reslice (bottom panels) views of *btd* RNAi embryos overexpressing Insc showing two classes of phenotype, visualized with 3xmScarlet-CaaX. White arrows, PMG; magenta arrows and magenta dashed outlines, head-trunk buckles; cyan arrows and cyan dashed outlines, small buckles near or inside MDs. Time is relative to the onset of gastrulation. Scale bars, 30 µm.

## Methods

### Experimental Animals

*Drosophila* embryos were collected on apple juice agar plates with yeast paste at 22°C or at temperatures indicated in the section below. The laboratory cultures of *Megaselia abdita* (Sander strain) and *Chironomus riparius* (Bergstrom strain) were maintained as described; *Megaselia* embryos were collected on apple juice agar plates with fish food paste at 25°C, *Chironomus* embryos as freshly deposited egg packages at ambient room temperature (23-26°C)^40^. The laboratory culture of *Clogmia albipunctata* was maintained as described; embryos were obtained after experimental egg activation through osmotic shock ^41,42^.

### *Drosophila* genetics and transgenic lines

*Drosophila* lines used for live imaging were *MyoII-eGFP* (also known as *Sqh-eGFP*)^43^, *Gap43-mCherry*^44^, *MyoII-mKate2*^45^, and *mat-tub-3xmScarlet-CaaX* (this study). The membrane imaging line *mat-tub-3xmScarlet-CaaX* was cloned into the pBabr vector containing the mat-tub promoter (a gift from D. St. Johnston, Gurdon Institute, UK)^46^ and the spaghetti-squash 3′ UTR, followed by ΨC31 site-directed integration into the attP2 or attP40 landing sites at WellGenetics, Inc. (Taipei, Taiwan).

*Drosophila* mutant alleles used were *eve^R13^* (FlyBase ID: FBal0003885), *btd^AX^*(FlyBase ID: FBal0030657), *stg^7M53^* (FlyBase ID: FBal0016176), and the quadruple mutant^25^ *knirps^IID48^* (FlyBase ID: FBal0005780) *hunchback^7M48^* (FlyBase ID: FBal0005395) *forkhead^E200^* (FlyBase ID: FBal0004007) *tailless^L10^* (FlyBase ID:FBal0016889). In live imaging experiments, the mutant embryos were identified based on the absence of a balancer-linked reporter construct, hb0.7-Venus-NLS, inserted on the FM7h, CyO, or TM3 balancer^9^.

To generate the *eve1^KO^* line, an *eve* genomic rescue construct, *eve^CH322–103K22^-mNeonGreen,* was first created using *P[acman]^CH322–103K22^* (BACPAC Resources, Center at Children’s Hospital Oakland Research Institute), a BAC construct that encompasses the entire *eve* locus, from which the stop codon of *eve* was replaced with mNeonGreen following a linker (N-ter-GSAGSAAGSGEV-C-ter) via a standard protocol ^47,48^. To completely eliminate *eve* expression in the Eve1 region, the stripe1 (st1; +6.6 kb to +7.4 kb relative to the transcriptional start site of *eve*)^19^ and late element (LE; –6.4 kb to –4.8 kb)^20^ enhancers were deleted from *eve^CH322–103K22^-mNeonGreen* through homologous recombination using the following homology arm sequences: st1– left: GCAAGTCCGAGACAAATCCACAAATATTGTCAACTCTTTGGCTCTAATCTG, right: CCAAGGCCGCAAAGTCAACAAGTCGGCAGCAAATTTCCCTTTGTCCGGCGA; LE – left: TTGCGTTTGAGCTACGTTACTTACATTTTTCCCACATGAGTCGGGCATACA, right: TCGATGGGTTGGTCACAATGTGGTGGCCTCTCAACATTGCAAGGCTCTTAC. The resultant BAC construct, *eve^CH322–103K22^-mNeonGreenΔst1ΔLE* (Fig. 2 Supplement 1a) was integrated into PBac{y[+]-attP-3B}VK00033 at Rainbow transgenics, USA, and crossed into the *eve^R13^* mutant line to generate the *eve1^KO^* line. Identification of *eve1^KO^*embryos in live imaging experiments were performed as above, based on the absence of a balancer-linked reporter construct, hb0.7-Venus-NLS, inserted on the CyO^9^.

For Insc overexpression, males of *UAS-insc* were crossed to *nos-GAL4-GCN4-bcd3’UTR*, which directs target gene expression in the head region of resultant embryos^49^. The flies were incubated at 25°C. For opto-DNRho1 experiments, females of *UASp-pmGFP-CIBN; UASp-CRY2-Rho1[N19, Y189]*^21^ (a gift from B. He, Dartmouth College, USA) were crossed to males of *matαTub-Gal4VP16^67C^; matαTub-Gal4VP16*^15^ double driver containing the *mat-tub-3xmScarlet-CaaX* imaging marker. The resultant F1 flies were used to set up egg deposition cages that were kept at 18°C for collection of embryos used in the experiments.

### Protein tree

Predicted protein sequences of *even-skipped* and *buttonhead* were used as query to identify closely related genes in *Drosophila* and putative orthologs in *Megaselia abdita* and *Chironomus riparius* using BLAST. Protein alignments were performed in Geneious by MUSCLE alignment with standard parameters. The protein tree was assembled using Jukes-Cantor as the genetic distance model and UPCMA tree build method, with a bootstrap of 1000 replicates.

### Cloning, and mRNA and dsRNA synthesis

*Cri-btd*, *Cri-eve*, *Cri-insc*, *Cri-sqh*, *Mab-btd*, and *Mab-eve* were identified from transcriptome sequences and cloned after PCR amplification from cDNA. In vivo reporters for cell outlines and MyoII in *Chironomus* were based on GAP43-linker-eGFP and Cri-Sqh-linker-eGFP fusion constructs, which were injected as *in vitro* synthesized mRNAs. The template of the Gap43-eGFP fusion construct was generated by in-frame Gibson cloning of the Gap43 encoding sequence, a short linker (GSAGSAAGSGEV), and a previously published pSP35T expression vector (pSP-Mab-bsg-eGFP) with 3’-terminal eGFP^50^. Analogously, the template of the Cri-Sqh-eGFP fusion construct was generated using a full length fragment of *Cri-sqh* amplified by PCR from cDNA. Nascent mRNAs were generated using SP6 polymerase, followed by capping and polyA-tailing with dedicated Capping and PolyA kits (Cellscript). Synthesized mRNA was dissolved in H_2_O.

For *btd* RNAi experiments in *Drosophila*, double stranded RNA (dsRNA) was synthesized on templates that contain the T7 promoter sequence (5’-TAATACGACTCACTATAGGGTACT-3’) at each end using a MEGAscript T7 kit (Ambion); templates were amplified from 0–4 h embryonic cDNA using specific primers (5’-AGCAGATGACGACGACAACA-3; 5’-TACTCGGACTTCATGTGGCA-3). For *insc* RNAi experiments in *Chironomus*, dsRNA was synthesized as previously described^51^. The dsRNAs comprised the following gene fragments (pos. 1 refers to first nucleotide in ORF): *btd*, pos. 1487 to 1817; *Cri-insc*, pos. 466 to 1892.

### Injections

For dsRNA injections in *Drosophila*, 0-1 hr old (up to stage 2) embryos were collected, dechorionated with bleach and mounted on an agar pad. The mounted embryos were then picked up using a coverslip painted with glue (prepared by immersing bits of scotch tape in heptane), desiccated for 10-14 min using Drierite (W. A. Hammond Drierite Co.) and covered with a mixture of Halocarbon oil 700 and 27 (Sigma-Aldrich) with a ratio of 3:1. Needles for injection were prepared from micro-capillaries (Drummond Microcaps, O.D. 0.97 mm, I.D. 0.7 mm) pulled with a Sutter P-97/IVF and beveled with a Narishige pipette beveller (EG-44). Injections were performed on a Zeiss Axio Observer D1 inverted microscope using a Narishige manipulator (MO-202U) and microinjector (IM300). A volume of ∼144 pL solution with a concentration of 1.1-1.6 μg/μl dsRNA was injected into the embryo. Embryos were kept at 25°C after injection in a moist chamber until early to mid-cellularization, followed by live imaging.

For injections in *Chironomus*, embryos were collected, prepared, and injected essentially as described^40^. Embryos were injected before the start of cellularization (approximately four hours after egg deposition), and then kept in a moist chamber until the onset of gastrulation. Throughout all procedures, embryos were kept at 25°C (± 1°C). Owing to their small size, *Chironomus* embryos (200 mm length) were always injected into the center of the yolk (50% of A-P axis). Embryos were injected with dsRNA typically at concentrations of 300 to 700 ng/ml; mRNA was injected typically at concentrations of 1.5-2.5 μg/μl (Cri-Gap43-eGFP and Cri-Sqh-eGFP). LiveAct-mCherry protein was injected at ∼4.5 mg/ml.

### Live imaging

Live imaging of *Drosophila* embryos was done using two-photon scanning microscopy with a 25X water immersion objective (N.A.= 1.05) on an upright Olympus FVMPE-4GDRS system with an InSight DeepSee pulsed IR Dual-Line laser system (Spectra Physics) or an inverted Olympus FVRS-F2SJ system equipped with a Maitai DeepSee and an InSight DeepSee pulsed IR laser systems (Spectra Physics), or with a Plan-Apochromat 25X oil immersion objective (N.A.= 0.8) on a Zeiss LSM980 inverted microscope equipped with a Chameleon laser (Coherent Int). For eGFP and Venus, a tunable laser line was tuned at 920 nm and 950 nm, respectively, while for mKate2, mCherry or mScarlet either a fixed line at 1040 nm on the upright system, or a tunable line tuned at 1100 nm on the inverted system was used. Three different general settings were used: 1) a z stack of ∼80 µm depth, image size of 539.5 x185.5 µm and a z step size of 2 µm with a 90 s time interval for an *en face* lateral view or ventral view of the whole embryo, 2) a z stack of ∼60 µm depth. Image size of 253.5 x152 µm and a z-step size of 1.5 µm with a 50 s time interval for an *en face* lateral view of the head domain, 3) a z stack of ∼40 µm depth, image dimension of 208.3 x152 µm and a z-step size of 1 µm with a 45 s time interval for the recording of cell division dynamics in the MDs of the head.

Live imaging of *Chironomus* embryos was performed on a Leica SP8 confocal using a 63x glycerol immersion objective (N.A.=1.30). Z-stacks of ∼25 µm depth were acquired at a z-step size of 1 µm and 90 sec time interval. All recordings were performed at 25°C.

Time-lapse imaging to visualize germband extension was performed on Nikon Eclipse-Ti microscope, in differential interference contrast (DIC) mode, using a 20x objective (N.A.=0.8) for *Drosophila*, *Megaselia*, *Chironomus*, and *Clogmia*, with 1 frame every min. All recordings were performed at 25°C.

### Optogenetics

The opto-DNRho1 system^21^ contains a dominant negative form of Rho1 (DNRho1) lacking the membrane localization signal and tagged with the photosensitive Cryptochrome 2 (CRY2), and a membrane-anchored N-terminal domain of the CIB1 protein (CIBN). Following blue light illumination, CRY2 undergoes a conformational change to bind to CIBN, thereby recruiting DNRho1 to the plasma membrane to inhibit cortical actomyosin contractility. Fly crosses, cages for egg deposition, and embryos prior to processing were kept in the dark. To prevent unwanted photoactivation, embryos were processed, staged and mounted in a dark room with the desk lamp and the brightfield light source on the stereo microscope both covered by a light red filter (#182, Lee filters, UK). Imaging was performed on an Olympus FVMPE-RS system with a 25X (N.A.=1.05) water immersion objective and with the 1040 nm fixed laser line from an InSight DeepSee pulsed IR Dual-Line laser system (Spectra Physics) for the membrane marker 3xmScarlet-CaaX. Photoactivation of the CRY2-CIBN system was achieved with a 405-nm diode laser set at 0.1% laser intensity, which produced 5.48 µW power measured at the sample position. This protocol has been first benchmarked on VF formation to confirm that it resulted in a complete blockage of apical constriction^21^. The laser was set at 0% for sham control. Two experimental designs were used: 1) For lateral imaging with unilateral photoactivation (Fig. 2g, h; Fig. 2 Supplement 1d; Fig. 2 Movie S3), an ROI of 28.15 x 197.05 µm centered on ‘the pre-CF domain’^52^ covering the entire region of CF initiation along the D-V circumference was illuminated for a scanning duration of 3 s, beginning at 16∼33 min prior to the onset of gastrulation and repeated every 90 s; 2) For ventral imaging with bilateral photoactivation (Fig. 3a, b; Fig. 3 Movie S1), two ROIs of 33.78 x 33.78 µm each covering one side of the CF was illuminated, for a scanning duration of 3 s, repeated every 180 s. Photoactivation began 18-30 min prior to the onset of gastrulation.

### Immunofluorescence and fixed imaging

For antibody staining, embryos were fixed by a heat-methanol method^53^ and immunostained with mouse monoclonal anti-Neurotactin (1:20, BP106, Developmental Studies Hybridoma Bank), rabbit polyclonal anti-Eve (1:500, gift from M. Biggin, Lawrence Berkeley National Laboratory, USA), and rat polyclonal anti-Btd (1: 500, gift from E. Wieschaus). Imaging was performed on a Leica SP8 system using a 20x (N.A.=0.75) multi-immersion objective with oil immersion with a total z depth of 60-90 µm and a z-step size of 1.04 µm.

For DNA staining, embryos were fixed by heat and devitellinized as described^54^, followed by staining with DRAQ5 (1:1000, 1 hr). Imaging was performed on a Leica SP8 system with a 20x glycerol objective (N.A.=0.75) for *Drosophila*, *Megaselia*, and *Clogmia*, and with a 63x glycerol objective (N.A.=1.3) for *Chironomus*, with a z-step size of 1 µm in a z-range that covers a half of the embryo.

For in situ hybridization, embryos were fixed by a heat-formaldehyde method^50^. Transcripts were detected histochemically or fluorescently as described^55^, using RNA probes for *Mab-btd* (comprising 1473 nts from +1 to 1473, with pos. +1 referring to first nucleotide in ORF), *Mab-eve* (comprising 984 nts, from position 365 to 996 of the putative CDS and 351 nts of 3’UTR), *Cri-btd* (comprising 831 nts from 476 to 1306), *Cri-eve* (comprising 892 nts from 30 to 921), and *Cri-insc* (comprising 1427 nts from 466 to 1892) labeled with either digoxigenin or fluorescein.

### Image processing and quantification

Images were processed, assembled into figures and converted into videos using FIJI, Affinity Designer, and Adobe Illustrator. Quantitative data were analyzed and processed by Excel and custom-made Python scripts using Numpy, Pandas, and SciPy libraries. Plots were generated in GraphPad Prism or with Python scripts using Matplotlib and Seaborn graphic libraries. The sections below provide brief descriptions of image processing and analysis procedures.

### Surface projection

For *en face* views, the FIJI plugin Local Z Projector^56^ was used to project a surface of interest from a 3D stack onto a 2D surface, taking into account the curvature of the embryo. The reference plane that represents the contour of the embryo surface was derived from a Gaussian-blurred image of the original 3D stack image with a σ value of 2∼4, followed by binarization with a customized threshold, which results in a smooth height map for z projection. For an optimal projection, three main parameters were considered, 1) median post filter for the height map, 2) ΔZ, and 3) offset for the maximum intensity projection. These parameters were standardized such that errors at the boundary of the embryo with the highest curvature and auto-fluorescence from the vitelline membrane closest to the topmost Z-slice were avoided. Whole embryo projections for embryos expressing Gap43-mCherry were done with ΔZ=0 and offset= 3 or 4, while for embryos expressing 3xmScarletCaaX, ΔZ=0 and offset=2 or 3 were used. For embryos expressing MyoII-GFP and MyoII-mKate2, ΔZ=3 and offset=4 or 5 were used.

### Re-slice along the z-axis

Re-slices were created using the re-slice tool in Fiji. A straight line approximately perpendicular to the CF or the head-trunk buckle was drawn and positioned between MD5 and MD9 to ensure the consistency of the slicing positioning along the D-V axis in all genotypes, except for the *stg* mutants where there is no cell division and the *khft* mutants where MD pattern is partially disrupted.

### Time annotation

Time-lapse images of laterally mounted *Drosophila* embryos were aligned temporally based on the fact that the initiation timings of CF, VF and PMG are concurrent, marking the onset of gastrulation^9^. For embryos imaged on the lateral side, the onset of PMG invagination was annotated based on a visual criteria of PMG surface flattening and the first major dorsal movement of the pole cells, with the exception of *khft* mutants in which the PMG was absent, and VF onset was used to define gastrulation onset. For embryos imaged on the ventral side, the first frame at which the cells in the VF move out of plane was set as gastrulation onset.

Time-lapse images of laterally positioned *Chironomus* embryos were aligned temporally based on the observation that similar to *Drosophila* the initiation timings of VF and PMG are concurrent, marking the onset of gastrulation. For laterally imaged embryos (PMG not visible), the onset of gastrulation was determined based on a visual criteria of first collective ventral-ward cell movement.

Fig. 4 Supplement 3 puts the dynamics of head divisions in *Drosophila* and *Chironomus* head ectoderm on a common timeline based on these time annotations.

### Measurement of CF or buckle depth

The depth of the CF or the buckles was measured at the time point where the invagination reaches the maximum depth. The segmented line tool in Fiji was used to trace the depth of the furrow starting from the vitelline membrane to the basal end of the epithelial cell at the CF or buckle tip.

### Particle Image Velocimetry (PIV)

PIV was performed on surface projections using the iterative PIV plugin in Fiji (https://sites.google.com/site/qingzongtseng/piv?authuser=0#h.khn09c6h1n39). Only the anterior half of the embryo was analyzed. Progressively decreasing Image interrogation window size of 88, 44 and 22 pixels was done with a search window of size 178, 88 and 44 respectively for each successive PIV iterations. The output of the 2nd iteration of PIV was used for the final vector plot. Time alignment of multiple embryos was based on gastrulation onset as described above. MD1 telophase onset was the one frame before the first MD1 cell undergoing cytokinesis. For the vector plot, vector coordinates were translated such that the x-axis origin was set at the CF or the buckle, hence representing the head-trunk boundary, while the y-axis origin represents the lateral midline. For each time point (gastrulation onset and MD1 telophase), vector fields from three successive time frames were averaged. Averaged vectors from multiple embryos were plotted with the size of the arrow representing the magnitude of the velocity and therefore the flow speed. The vector lengths were extended by a factor of two for better visualization.

### Nuclear density

Nuclear density was assessed at the end of cellularization by nuclear segmentation using ‘Find Maxima’ in Fiji, followed by manual corrections and Voronoi tessellation to generate a pseudo-cell territory around each maximum; territories at the edge of the image were excluded. The median of the pseudo-cell territory area was used to estimate nuclear density.

### Measurement of surface area in mitotic cells and regions

The surface area of individual mitotic cells or mitotically active regions was obtained by surface area segmentation of embryos expressing a fluorescent reporter for membranes using the TissueAnalyzer^57^ plugin in Fiji. *Chironomus* embryos expressed Gap43-eGFP, and segmentation was performed on a single z-slice 4 µm below the cell apex; *Drosophila* embryos expressed 3xmScarlet-CaaX, and segmentation was performed on surface projections generated using the FIJI plugin Local Z Projector as described above. For the analysis of individual dividing cells, segmentation was carried out separately for each timepoint and cells were then tracked manually over time (24 out-of-plane and 18 in-plane dividing cells from 3 embryos). For each time point, the area of tracked cells was extracted. Following an in-plane division, both daughter cells were tracked and their area summed. Following an out-of-plane division, only the cell remaining at the surface was tracked; occasional short contacts of the bottom cell to the apical surface were not included in the analysis. Time annotation was performed respective to each individual cell’s onset of telophase. For the quantification of mitotically active domains, the largest possible representative area of a given mitotic domain with fully traceable cells was chosen. Domain areas were tracked and extracted manually for 3 embryos. For domain analysis time annotation was based on onset of gastrulation (described above).

### Analysis of ventral midline deviation

For an estimate of deviation from the expected linear ventral midline, the actual ventral midlines were marked manually using the segmented line tool in FIJI at the onset of gastrulation (‘onset’) and a later time point (‘mid’) when the midline exhibits maximal distortion. The resultant ROIs were converted into points using built-in macro function Roi.getContainedPoints in FIJI, and the coordinates were analyzed using a Python code. To compute deviation, a linear fit to the midline coordinate was first generated as the expected midline for the onset stage, which was then translated to the linear fit for the midline coordinates marked at the ‘mid’ stage, while preserving the slope, as the expected midline for the mid stage. For each stage, the mean distance from the marked midline coordinates to the expected midline was calculated and plotted as ‘Deviation’ in Fig. 3c, f.

### Statistical Analyses

Python scripts using SciPy library were implemented to conduct one-way ANOVA followed by Tukey’s multiple comparison post hoc test for comparing means from more than two groups, and Mann-Whitney U test as a non-parametric independent test. All of the statistical details of experiments, including the number of experiments (n), which represents the number of embryos used unless otherwise noted, were indicated in the figure legends.

Statistical analysis to compare the blastoderm cell densities across species was done using GraphPad Prism, where we estimated the statistical significance using one-way ANOVA with Kruskal-Wallis non-parametric test, without correcting for multiple comparisons (Uncorrected Dunn’s test). The same program was also used to calculate medians and the 95% confidence intervals on the median.

## Data and Code Availability

Data and codes developed for data analysis are available upon request from the corresponding authors.

## Acknowledgements

We thank the BACPAC Resources, Center at Children’s Hospital Oakland Research Institute, the Developmental Studies Hybridoma Bank, the Bloomington and Kyoto *Drosophila* Stock Centers, Yohanns Bellaïche, Mark Biggins, Stefano De Renzis, Nathalie Dostatni, Bing He, Shigeo Hayashi and Eric Wieschaus for sharing reagents; Shigeo Hayashi, Lars Hufnagel, and Jochen Wittbrodt for support; Anthony Eritano for initial characterization of the *eve1^KO^* line; Lucas Schütz and Atalay Tok for initial conceptualization and observations of division orientation in *Chironomus*; Laura Popp and Maike Fath for cloning and initial characterization of gene expression in *Chironomus*; members of the Hayashi, Obata and Yoo laboratories for discussions; Anna Erzberger and Girish Deshpande for critical reading and comments on the manuscript. This work was supported by the core funding at RIKEN BDR; the Human Frontier Scientific Program (HFSP) Young Investigators grant to S.L. and Y.-C.W. (RGY0082/2015); a Research Grant by the Deutsche Forschungsgemeinschaft to S.L. (LE 2787/3-1); G.K. was supported by HFSP Long-term postdoctoral fellowship (LT000597/2019-L).

## Author Contributions

Y.-C.W and S.L. conceived, designed and supervised the study; B.D., V.K., G.K, M.S., M.T., and Y.-C.W. generated the reagents, wrote software code, and performed the experiments; B.D., V.K., G.K, M.S., M.T., Y.-C.W., and S.L. analyzed the data and prepared the visualization; B.D., V.K., G.K, Y.-C.W and S.L. wrote and edited the manuscript.

## Additional Information

Supplementary Information is available for this paper. Reprints and permissions information is available on the journal website. Correspondence and requests for materials should be addressed to Y.-C.W. and S.L.

## Footnotes

1 This study has been submitted back-to-back with the present manuscript. Both pre-prints are available on bioRxiv.

